# Obtaining tertiary protein structures by the ab-initio interpretation of small angle X-ray scattering data

**DOI:** 10.1101/572057

**Authors:** Christopher Prior, Owen R Davies, Daniel Bruce, Ehmke Pohl

## Abstract

Small angle X-ray scattering (SAXS) has become an important tool to investigate the structure of proteins in solution. In this paper we present a novel ab-initio method to represent polypeptide chains as discrete curves that can be used to derive a meaningful three-dimensional model from **only** the primary sequence and experimental SAXS data. High resolution crystal structures were used to generate probability density functions for each of the common secondary structural elements found in proteins. These are used to place realistic restraints on the model curve’s geometry. To evaluate the quality of potential models and demonstrate the efficacy of this novel technique we developed a new statistic to compare the entangled geometry of two open curves, based on mathematical techniques from knot theory. The chain model is coupled with a novel explicit hydration shell model in order derive physically meaningful 3D models by optimizing configurations against experimental SAXS data using a monte-caro based algorithm. We show that the combination of our ab-initio method with spatial restraints based on contact predictions successfully derives a biologically plausible model of the coiled–coil component of the human synaptonemal complex central element protein.

**SIGNIFICANCE:** Small-angle X-ray scattering allows for structure determination of biological macromolecules and their complexes in aqueous solution. Using a discrete curve representation of the polypeptide chain and combining it with empirically determined constraints and a realistic solvent model we are now able to derive realistic ab-initio 3-dimensional models from BioSAXS data. The method only require a primary sequence and the scattering data form the user.

## INTRODUCTION

Biological small angle X-ray scattering (BioSAXS) is an increasingly important method for characterising protein structures in solution (1–3). Its primary advantages over complmentary techniques such as crystallography and NMR is its ability to provide information under native conditions about large protein molecules not accessible by complementary methods. However, there is a price to pay for this advantage; the random motion and orientation of molecules in solution leads to a loss of information due to an effective averaging of the scattering, leaving only information about the protein’s intra-molecular distances not their spatial orientations (4). Furthermore, scattering extends only to low resolution providing limited experimental data. The correct interpretation leading to meaningful biological results remains therefore challenging (5).

Two main methods have been developed to interpret BioSAXS data. The first assumes an accurate 3D model of the protein backbone, usually derived from X-ray crystallography (6–9). This model is used to calculate the X-ray scattering curve once the excluded solvent volume is taken into account. A major advance, first presented in the CRYSOL algorithm (6), was the inclusion of the solvation layer - the ordered water molecules at the surface of the protein. CRYSOL as well as the FOXS package, developed by Schneidmann-Duhovny *et al* (7), adjust an implicit “shell” of scattering (implicit meaning they do not model individual solvent molecules). Other packages treat the shell explicitly using either molecular dynamics (AquaSAXS) (8) or a geometric filling approach (the SCT suite) (9). Allowing for a shell which can have gaps and fill cavities in the protein model gives a more reliable fit to the data (5). An extension of this approach is to use all atomistic modeling with PDB structures as a start point (10, 11), the application of such techniques, however, can require signification technical expertise. The second method does not assume an initial structure (ab-initio) but simplifies the protein model as either a volume (12) or a chain (13) of scattering beads without explicit secondary structure. These methods are hence applicable to *de novo* structural prediction, but the lack of secondary structure means interpreting these predictions is a difficult task (5).

Here we propose an alternative ab-initio technique which uses a curve model of the 3D structure of the polypeptide chain, this description has a much reduced number of parameters by comparison to all atomistic models. Similar curve models have been previously proposed (14–16) but not for the purpose of interpreting BioSAXS data. The model is parameterised by consecutive discretised descriptions of the four major secondary structural elements, *α*-helices, *β*-strands, flexible sections and random coils. The permissible geometry of these curves is restricted by empirically determined constraints, which are akin to Ramachandran constraints (17). To use the model for interpretation of BioSAXS data the polypeptide chain model is combined with a water model for the first hydration shell and an empirically calibrated scattering model. The geometry of the model can then be optimized against the experimental BioSAXS data, a critical factor, novel to our curve representation of the polypeptide chain, is the construction of empirical probability distributions for the model parameters. These distributions serve the dual purpose of preferencing commonly observed secondary structures in the set of potential chain models, whilst simultaneously allowing for predictions with rare/novel but physically permissible secondary structure. An advantage of this technique ab-initio interpretation of BioSAXS data, by comparison to the established bead models (12, 13), is that by accurately characterizing the protein’s secondary structure it can reliably incorporate additional structural information in order to improve the results of the technique. In this study contact predictions, based on sequence alignments alone, are used to improve the model predictions. A final advantage of the code developed is that its only input requirements are the primary sequence and scattering data, so places only basic technical requirements on the user for its use.

We first applied this new methodology to data of well characterized model protein Lysozyme before moving to the BioSAXS data of structural core of the human synaptonemal complex central element protein 1 (SYCE1). This protein represents an essential structural component of the synaptonemal complex (SC) that binds together homologous chromosomes during meiosis and provides the necessary three-dimensional environment for crossover formation (18–20). The SC is formed of oligomeric *α*-helical coiled-coil proteins that undergo self-assembly to create a lattice-like assembly (21–23). In a recent biochemical and biophysical study, human SYCE1 was shown to adopt a homodimeric structure in which its structural core is provided by residues 25-179 forming an anti-parallel coiled-coil (24). Further, the structural core was expressed in an engineered construct in which two SYCE1 25-179 sequences were tethered together through a short linker sequence (GQTNPG). This construct faithfully reproduced the native structure, and substantially improved protein stability in solution (24). In this study, using secondary structure predictions and distance restraints purely based on the sequence of the protein alone, an excellent model of an anti-parallel extended but bent coiled-coil is derived, which is fully consistent with biological data.

## METHODS

First we describe the reduced parameter protein model we use to interpret the BioSAXS data. This is composed of a polypetide chain curve model with a surrounding explict hydration shell. Empirically calibrated structure factor functions for each constituent element of the model are constructed to produce theoretical scattering curves for this tertiary structure model.

### Polypeptide chain

The polypeptide chain is represented as a set of points in 3D space 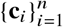, the positions of the C^*α*^ atoms in each amino acid. The geometry of four consecutive points (**c**_*i*_, **c**_*i*+1_, **c**_*i*+2_, **c**_*i*+3_) can be characterized by two parameters, the curvature *κ* and torsion *τ*. *κ* is defined by the unique sphere made by the centre of the joining edges (see Figure 1(a)), the smaller the sphere the more tightly the curve joining the points fold on themselves, *τ* measures the chirality of the section, it is positive for right-handed coiling negative if left-handed. More precise definitions are as follows:

**Figure 1:**
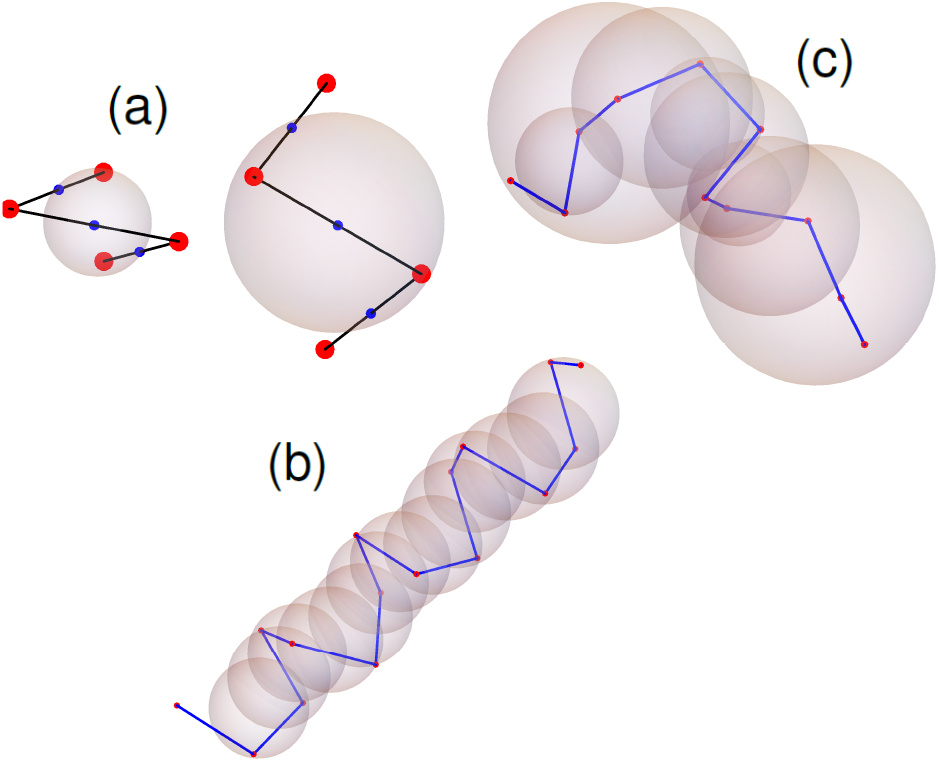
Figures depicting elements of the backbone model. (a) curve subsections (**c**_*i*_, **c**_*i*+1_, **c**_*i*+2_, **c**_*i*+3_) (red points) and their mid section points (**c**_*m*1_, **c**_*m*2_, **c**_*m*3_) (blue), the first example is more tightly wound and has a smaller sphere, hence a higher *κ* value. The sphere defined by these mid-section points is shown, the inverse of it’s radius is the curvature *κ*. (b) an *α*-helical section with uniformly similar (*κ*, *τ*) values. (c) a flexible (linker) section with varying (*κ*, *τ*) values.

### Curvature *κ*

A section of four residues defined by the points (**c**_*i*_, **c**_*i*+1_, **c**_*i*+2_, **c**_*i*+3_) defines three edges with midpoints **c**_*ml*_ + (**c**_*i*+*l*−1_ + **c**_*i*+*l*_)/2, which in turn define the curvature sphere (25) (see Figure 1). The curvature, the inverse of its radius is

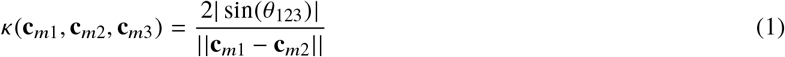

where *θ*_123_ is the angle between the vectors **c**_*m*1_ − **c**_*m*3_ and **c**_*m*2_ − **c**_*m*3_.

### Torsion *τ*

Three points define a plane (with unit normal vector **n**) and the four points (**c**_*i*_, **c**_*i*+1_, **c**_*i*+2_, **c**_*i*+3_) define two planes through their unit normal vectors **n**_2_ and **n**_2_ respectively:

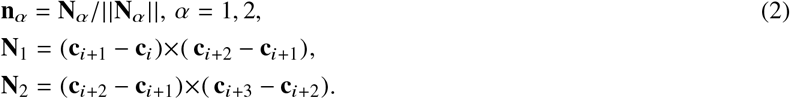

The torsion is the (length weighted) angle these planes make with each other,

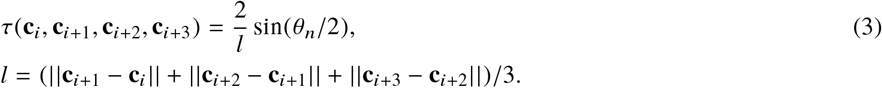

with *θ*_*n*_ is the angle between **n**_1_ and **n**_2_, see *e.g.* (26).

The algorithm for generating a curve of length *n* from *n* − 3 pairs of values of (*κ*_*i*_, *τ*_*i*_) is as follows: Consider a section of curve of length *m* and *m* − 3 pairs (*κ*_*i*_, *τ*_*i*_), whose three initial points **c**_1_, **c**_2_, **c**_3_ are randomly chosen (with fixed separation distance *R* = 3.8). Since scattering expressions are invariant under an arbitrary translation and rotation ((4)) the exact values of the first two points do not matter (as long as their separation is *R*). The third point is a structural degree of freedom but it is restricted such that the C^*α*^-C^*α*^ distance between **c**_1_ and **c**_3_ is greater than *R*. Once these points are specified the fourth point will be

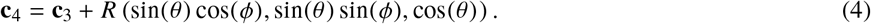

with *θ* = ∈ [0, *π*], *ϕ* ∈ [0, 2*π*]. The set (**c**_1_, **c**_2_, **c**_3_, *θ*, *ϕ*) define four points and hence *κ* and *τ* values. Using values of *κ*_1_ and *τ*_1_ equations (1) and (3) are solved for *θ* and *ϕ*, this gives **c**_4_. The next point **c**_5_ can similarly be found from the values *κ*_2_ and *τ*_3_, and so on until all *m* − 3 (*κ*_*i*_, *τ*_*i*_) have been used to yield the *m* points **c**_*i*_. Examples of an alpha-helical and flexible linker sections (taken from the structure of Bovine serum albumin (PDB=3V03) (27)) are shown in Figure 1(b) and (c).

### Secondary structure geometry restraints

In order to derive geometric constraints C^*α*^ coordinates were extracted from over from a set of over 60 protein structures for which high-resolution crystal structures are available in the Protein Data Base (PDB) and the *κ* and *τ* values calculated for all sub-sections (**c**_*i*_, **c**_*i*+1_, **c**_*i*+2_, **c**_*i*+3_). The *κ*-*τ* pairs are shown in Figure 2(a). There are three main populations of values (preferential regions). As shown in section 1.3 of the supplementary material these regions of (*κ*, *τ*) space correspond to the three preferential domains of Ramachandran space ((17)). Using the PDB’s secondary structure annotation this data was split into categories of *β*-strands, *α*-helices and the rest which are not identified (referred to here as linkers). To account for random coils the data were further divided into subsets whose values remained in one preferential domain (as in Figure 1(b)) and those whose *κ*-*τ* values belong to multiple domains (like Figure 1(c)). For each set of data a representative probability density function (P.D.F.) was calculated using Kernel smoothing techniques (28) (for details see sections 1.4 and 1.5 of the supplementary material), an example is shown in Figure 2(b).

**Figure 2:**
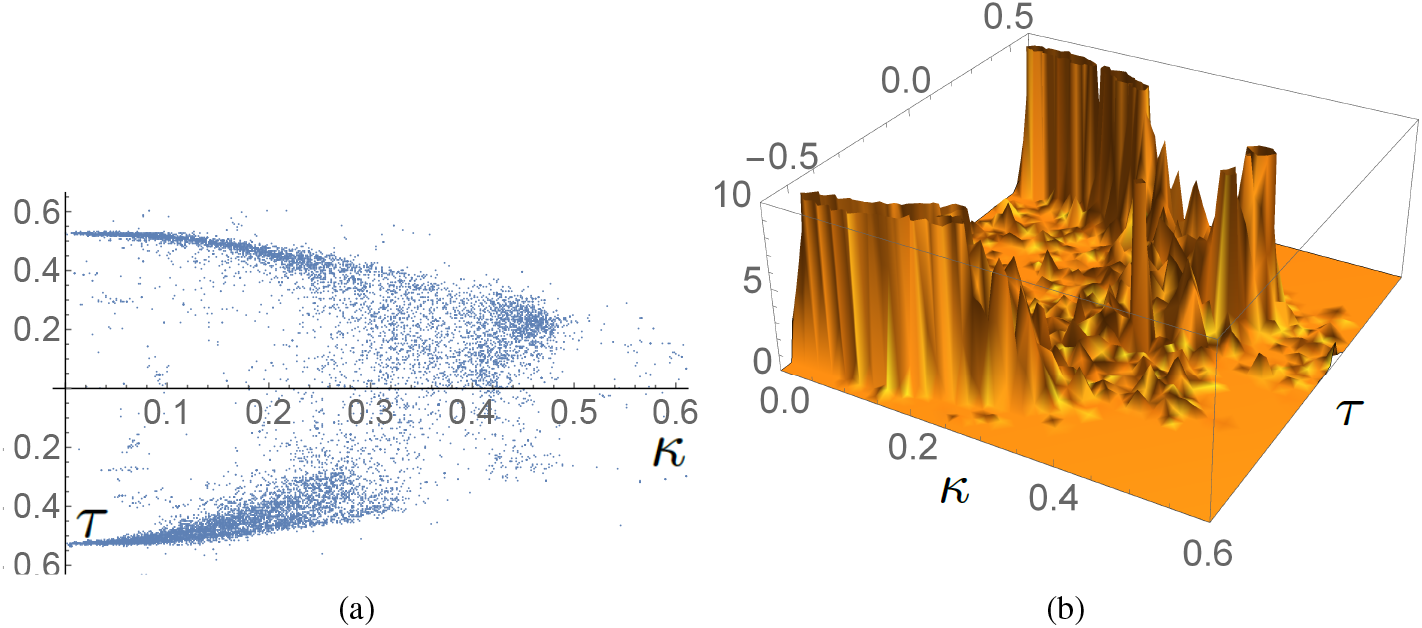
Illustrations of the *κ*-*τ* spaces used to impose realistic geometry constraints on the polypeptide chain. (a) (*κ*, *τ*) pairs obtained from crystal structures, plotted as points with *κ* on the horizontal axis and *τ* the vertical axis. (b) is a P.D.F, created from the data in (a), which correspond to linker sections. There are three distinct domains of high probability corresponding to the preferred corresponding to the preferred secondary structural elements.

### Generating models from secondary structure annotation

In order to generate models based on secondary structure information alone a protein of *n* amino acids is split into *l* distinct sub-domains of length 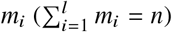. Each section *l* is classified as *α*-helical, *β*-strand or linker, for the purpose of testing and calibration the PDB file’s secondary structure assignment was used to perform this task. For each section of length *m*_*i*_, *m*_*i*_ − 3 (*κ*, *τ*) pairs are drawn from an appropriate P.D.F. and the section is constructed. This process creates the *l* individual secondary structures, which must then be linked together. Two neighboring sections with specified geometry (for example an *α* helix and linker) still have a relative rotational degree of freedom. To ensure this remains physically realistic the geometry of the last three and first C^*α*^ positions of neighboring secondary sections were extracted from the PDB set and further PDF’s for the set of permissible (*κ*, *τ*) pairs of these joining sections were generated for each type of join (*i.e. α*-helix to linker or linker to *β* strand). So the final step of the process is to obtain all (*κ*, *τ*) values for the joint geometry and then construct the whole backbone. A precise mathematical description of this algorithm, *constrained backbone algorithm* (**CB**), is given in section 1.6 of the supplement. One example of a structure generated using this algorithm is shown in Figure 6(b), this particular structure was used as a starting point for an ab-initio structure optimization in this study.

### The hydration layer

Once the curve representation is obtained it is crucial to include a model of the hydration layer in order to generate realistic scattering curves. To this aim solvent molecules are placed in-between a pair of cylindrical surfaces surrounding the axis of a section of the backbone (Figure 3(a)). This layer is then reduced by removing all overlapping solvent molecules. This ensures the shell remains in hollow sections between the fold and on the protein surface, whilst the water molecules are removed form significantly folded regions. This method is illustrated in Figure 3 where the two cylinders of radius *R*_*c*_ (core) and *R_o_* (outer), *R*_*o*_ > *R*_*c*_ are centered on a section *i*’s helical axis (a). Consider a solvent molecule belonging to another section *j* whose nearest distance from the axis of section *i* is *R*_*s*_. If *R_s_* < *R*_*c*_ the solvent is too close to the backbone and removed. If *R*_*c*_ < *R*_*s*_ < *R*_*o*_ the solvent is classed as being shared by the sections *i* and *j* and only counted once.

**Figure 3:**
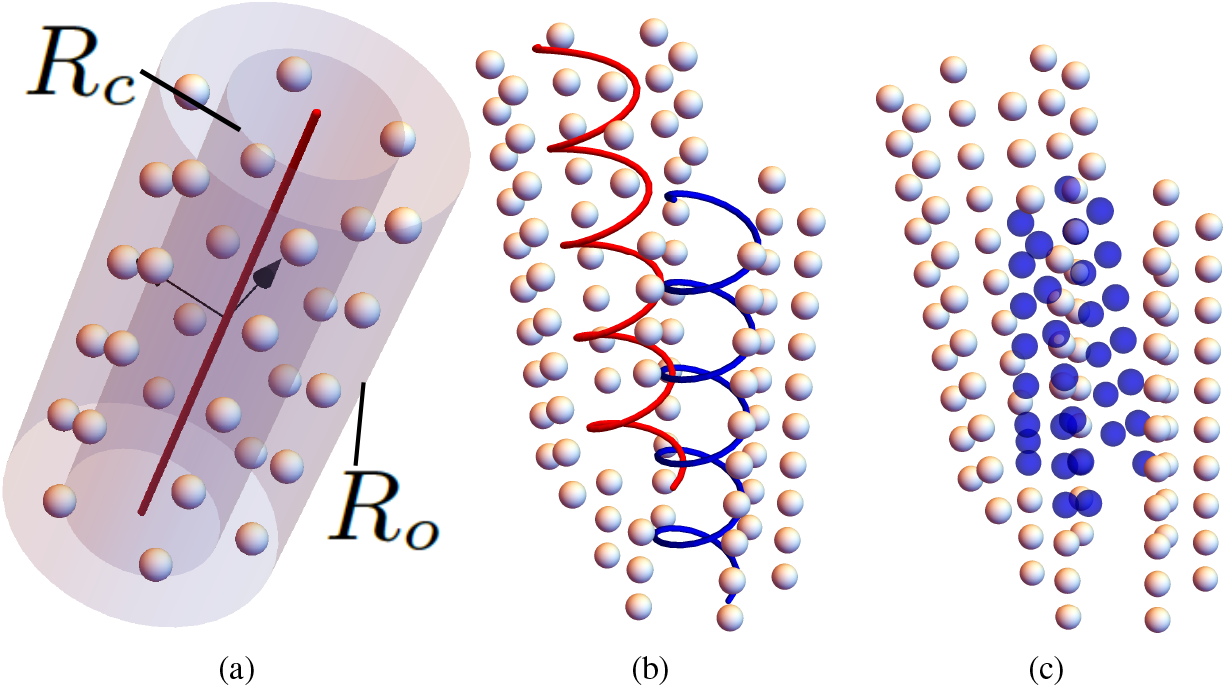
Visualizations of the hydration layer model. (a) the initial solvent layer, shown as silver spheres with the core *R*_*c*_ and outer *R*_*o*_ cylinders surrounding the axis of the section (red curve). (b) overlapping sections and solvent layers, (c) shows, in blue, the removed solvent molecules of the pair of sections shown in (b).

This process is applied to all solvent molecules from section *i* and *j* on each other, an example of the outcome is shown in Figures 3(b) and (c). Applying this process pairwise to all sections of a C^*α*^ backbone yields the final hydration layer.

The exact mathematical description of this hydration layer is detailed in sections 2.1-2.3 of the supplement. The values of the radii (*R*_*c*_, *R*_*o*_) and a number of other parameters controlling the solvent density were determined by fitting the model to high resolution crystal structures which contained the first hydration shell. An example model shell, generated with these parameters, is shown in comparison to the model solvent positions from the subatomic resolution structure of a phosphate binding protein from the PDB 4F1V (29) in Figure 4. It is shown the two distributions are statistically similar in section 2.4 of the supplement and hence that the model is a realistic representation of the average positions of the inner hydration shell.

**Figure 4:**
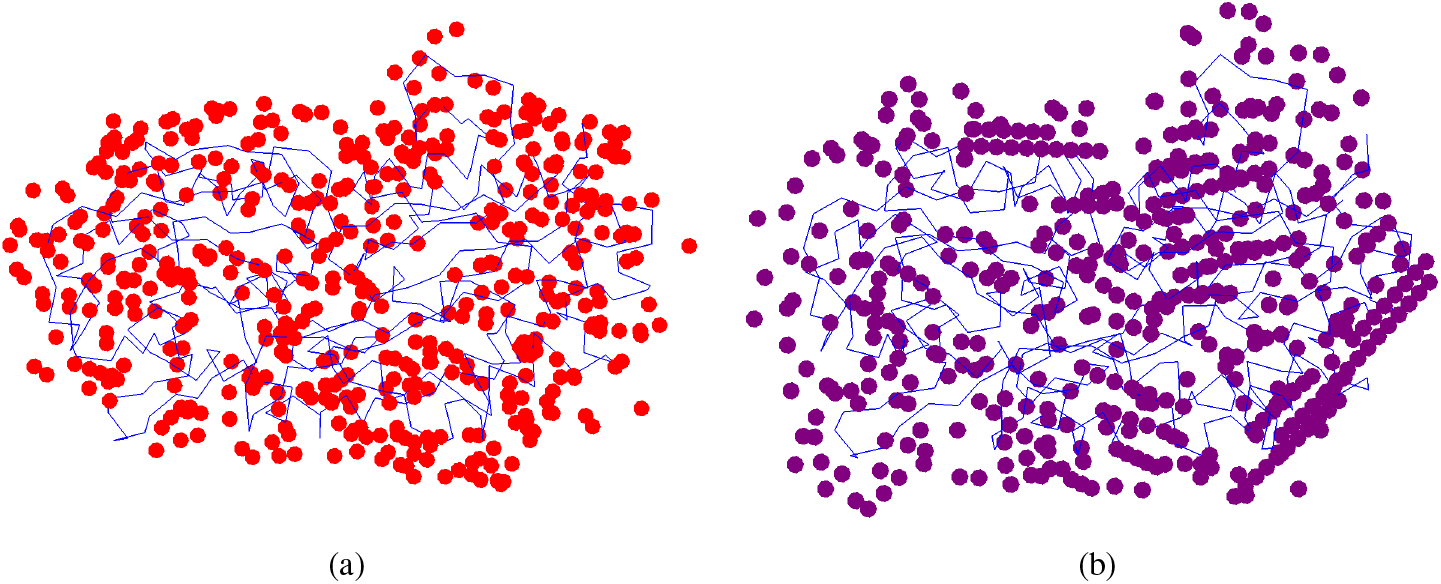
Comparisons of crystallographic and model solvent positions from the crystal structure of a phosphate binding protein PDB=4F1V, determined at an ultra- high resolution of 0.88 Å (29). (a) the PDB backbone and the relevant solvent molecules. (b) the model solvent positions (surrounding the same curve as in (a)) obtained with the experimentally determined hydration shell model parameters.

### The scattering formula

Once the polypeptide chain and hydration layer models are determined, the Debye formula (30),

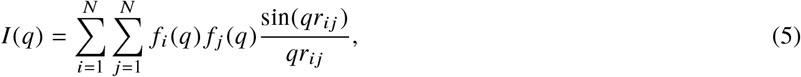

is used to calculate the scattered intensity *I* (*q*) as a function of momentum transfer *q* = *π* sin(*θ*)/*λ*. Here *N* is total number of C^*α*^’s and solvent molecules and *f*_*i*_ (*q*) the form factor for residue *i*. There are two types, one for an amino acid with an excluded volume correction and one for a solvent molecule which are defined as follows:

### Amino acid form factors

The form factor *f*_*am*_ of an amino acid, centered on the C^*α*^ atom position, are

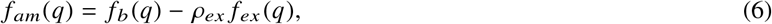

where *f*_*b*_ is the scattering of the amino acid in a vacuum and *f*_*ex*_ is the adjustment due to the excluded volume of solvent and *ρ*_*ex*_ a constant. Each amino acid is assigned the same scattering function *f*_*b*_(*q*), a five-factor exponential representation

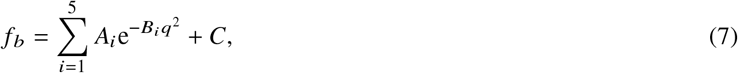

where 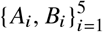 and *C* are empirically determined constants (a standard form used to fit molecular form factors (31)). The excluded volume effect is captured using an exponential model in the form

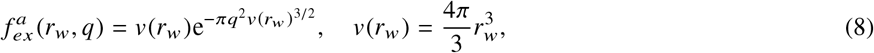

where *r*_*w*_ is the average atomic radius of the atom (6, 7, 13). To calculate the excluded volume for amino acids coordinates for all 20 amino acids (32), and values of *r*_*w*_ for Carbon, Nitrogen, Oxygen, Hydrogen and Sulphur (*e.g.* (33)) were used to compute the excluded volume scattering, centered at the C^*α*^, through

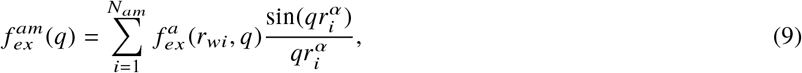

where 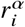 is the distance of atom *i* from the C^*α*^ molecule and *N*_*am*_ the number of atoms in the amino acid. Since *f*_*b*_ does not discriminate individual amino acids this value 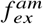 was averaged over all 20 amino acids, weighted by their abundance in globular proteins (see (34)). This averaged function, shown in Figure 5(e), gives *f*_*ex*_(*q*). Finally (6) includes a constant *ρ*_*ex*_ which modulates the effect of the excluded volume scatter by comparison to *f*_*b*_, this value is constrained to lie within 0.75 and 1.25 (similar constraints are used in (6, 7, 13)). The scattering form for an individual water molecule in the hydration layer is

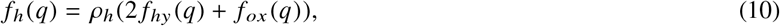

where *f*_*hy*_ and *f*_*ox*_ are the vacuum scattering of Hydrogen and Oxygen respectively (31). The constant *ρ*_*h*_ was empirically determined (as in (7)). A detailed description of the parameter determination method is given in section 3 of the supplementray materials.

**Figure 5:**
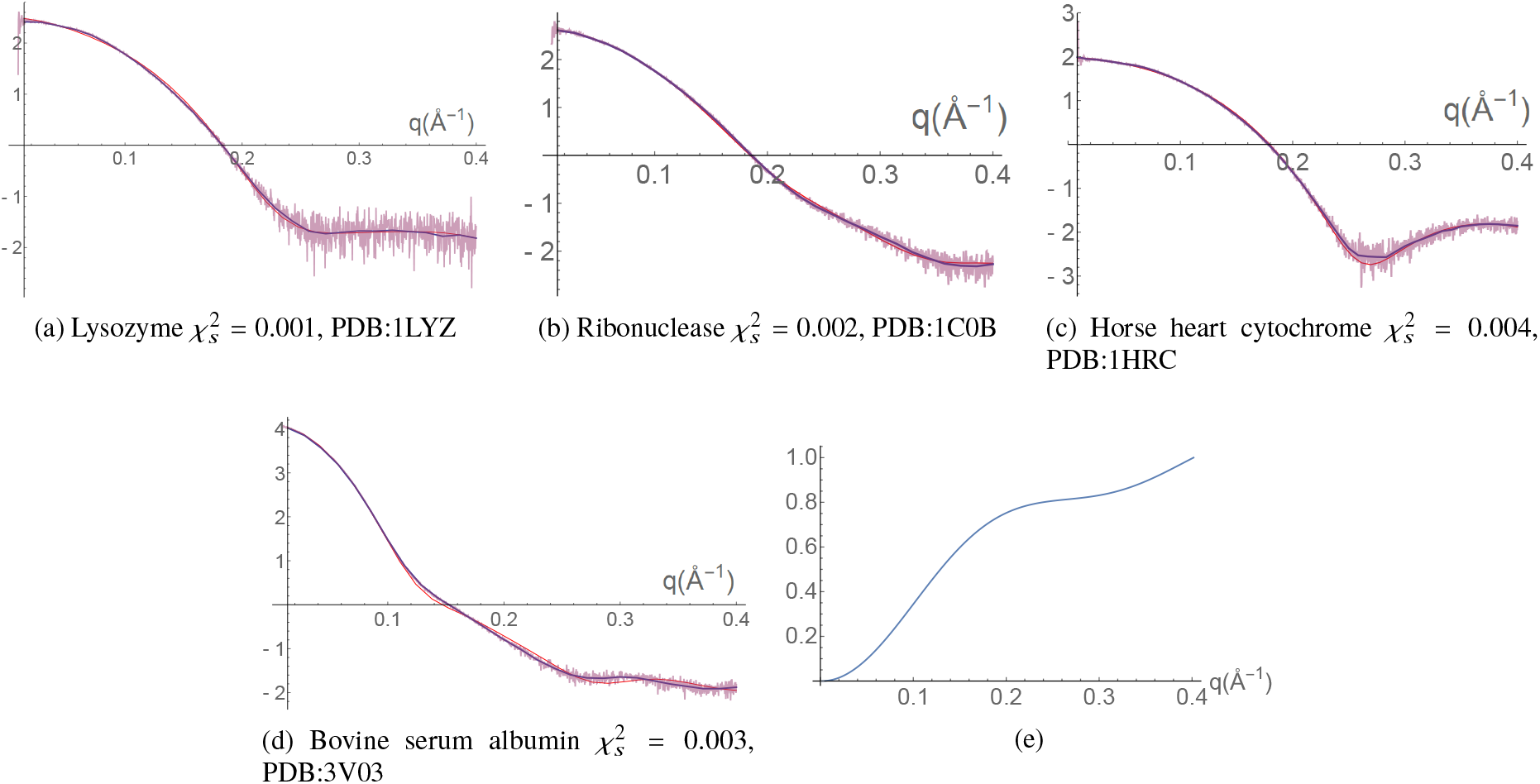
Fits to scattering data for various molecules using appropriate C^*α*^ coordinates as a backbone model 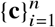 (see chapter 3 of the supplementary notes for details). In panels (a)-(d) The data scattering data is shown overlayed by the smoothed data used for fitting (blue curve) and and the model fit (red curve). Panel is the (e) the averaged scattering function 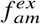 obtained by averaging the scattering parameters obtained from (a)-(d).

These form factors require a number of constants to be specified. To determine these constants the hydration layer model defined the previous section was applied to backbone models obtained from X-ray structures with high quality BioSAXS data available from the SAS database (35). The parameters were then obtained by a global optimization to the experimental data in which (5) was applied to this composite backbone-hydration shell model (see section 3 of the supplement). The quality of fit obtained, as shown in Figure 5, is a further endorsement of the hydration layer model introduced above.

### Evaluating structural similarity

In the next step the geometry of each model generated by the CB algorithm is optimized by refinement against the scattering data. However, since the problem is under-determined, many models will fit the experimental data so a method is required to compare structures and determine which predictions are “essentially the same” in that they only differ by small local conformational changes (as one should expect in solution). The standard methods in protein crystallography for comparing similar protein structures are based on root mean squared deviations (RMSD) where two structures are superimposed to minimize the sum of all distances of equivalent paired atoms (36, 37). This measure and variants on it are known to be overly sensitive to large deviations in single loops (as discussed in (36)). Unlike homologous crystal structures, which will often only differ by the change in a small subsection of the whole structure, the comparison here will be made between structures generated by a random algorithm, so the significant build up of relatively small individual RMSD errors is likely. In section 2.1 of the supplement a number of additional problems with using the RMSD measure in this context are discussed in detail. To mitigate these problems a novel and more robust approach based on knot theoretic techniques was developed.

### Knot fingerprints

Techniques from knot theory have previously been applied to identify specific (knotted) entanglements in protein structures (38). To compare two protein structures using knot theory the N and C termini need to be joined (39). As in (38) the procedure used here is to surround the backbone with a sphere, then choose two random points on the sphere and join the end termini to these points, finally this extended curve is closed with a geodesic arc. The knot is then classified (*e.g.* via Jones polynomials). This procedure is repeated a significant number of times (10000 in this study) and the most common knot (MCK) chosen to indicate the knotting of the curve. To obtain additional information the MCK is calculated for all subsets {**c**_*i*_ | *i* = *k*, *k* + 1 … *j*, *j* > *k*, *j* − *k* > 3} of the curve. One can then plot this data on a “staircase” diagram with *j* and *k* on the axes and each square of the domain colored by its most common knot (*e.g.* (40)) (examples of staircase diagrams are shown in Figure 6(c), (d) and (e)). The fingerprint is found to be preserved across protein families (40), even when there is low sequence identity (41).

**Figure 6:**
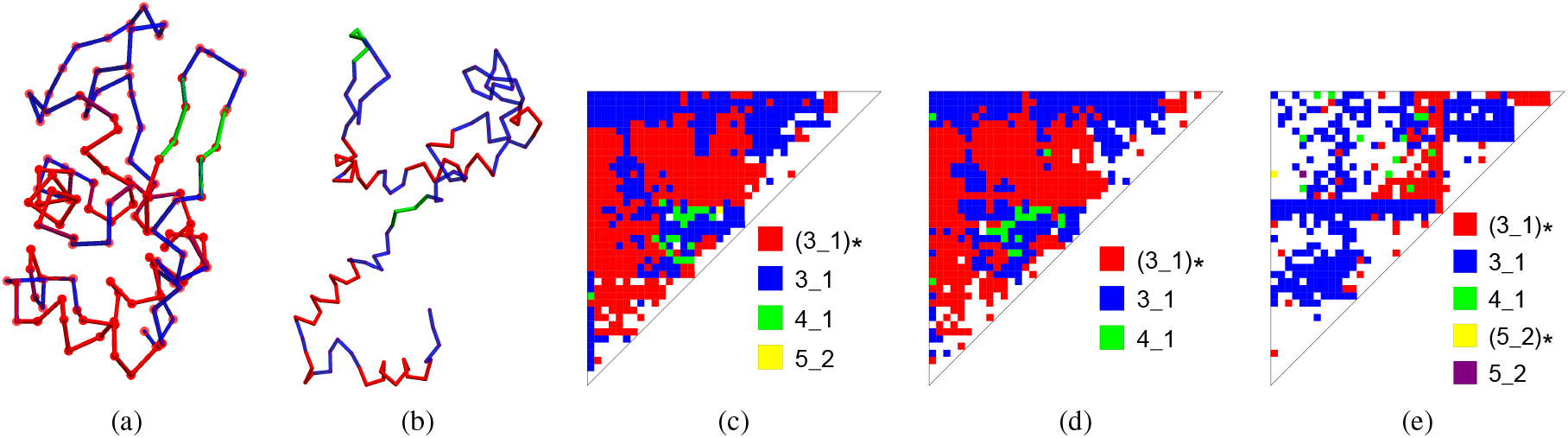
Secondary knot fingerprint analysis of the Lysozyme structure. (a) The C^*α*^ trace of Lysozyme (PDB 1LYZ (42)). The *α*-helices are shown in red, *β*-strand structures green, and linker sections light blue. (b) A random structure generated using the CB algorithm which has the same secondary structural elements as Lysozyme. This could be a starting model for the fitting procedure. Panels (c) and (d) are secondary fingerprints of two different crystal structure of Lysozome (1LYZ and the 1AKI respectively). The knot types are indicated (Rolfsen classification (43)), white spaces indicate no secondary knots (all knots were of the primary type). (e) Secondary fingerprint for the random structure shown in (b), it differs significantly from (c) and (d) and has a larger range of knots present.

### Secondary knot fingerprints

Figure 6(c) is the knot fingerprint for one set of Lysozyme coordinates (shown in Figure 6(a)), of the **second** most common knot identified during the random closure process. The secondary fingerprint shown in 6(d) is from a second set of Lysozyme coordinates, (c) and (d) are significantly similar. The secondary fingerprint (e) is derived from a CB generated backbone model, shown in (b), which has the same secondary structure sequence as the 1LYZ PDB. The secondary fingerprint differences between the correct structure (c) and the randomly generated structure (d) is immediately obvious. All primary (MCK) fingerprints in these cases are *identical* and all have the unknot as the MCK. It is clear secondary (and possibly Tertiary) knot fingerprints can differentiate un-knotted folds. A knot fingerprint statistic 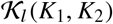 is defined in section 4.2 of the supplement which quantifies the weighted similarity of knot fingerprints at level *l* associated with the curves *K*_1_ and *K*_2_ (*l* = 2 for Figures 6(c)-(e)); it yields a value between 0, completely dissimilar, and 1, identically folded.

In section 4.3 of the supplement it is demonstrated that the statistic has the following properties. Firstly it quantifies crystal structures of the same molecule as highly similar 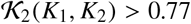 and randomly generated structures (with the same secondary structure sequence) as significantly dissimilar, generally 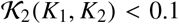 (see Figure 7(a)). Secondly it judges crystal monomer structures of similar length as being significantly different (typically 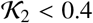), *i.e.* it can differentiate folds. Thirdly it is shown to have excellent properties under deformation. To demonstrate, *n* randomly distributed changes were applied to a crystal structure *K*_*pdb*_ using the CB algorithm. For each *n* 50 such structures *K*_*n*_ were generated and the values of the statistic 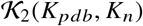 calculated. The results are plotted as a function of *n* in Figure 7(b) for Lysozyme. The mean value drops off rapidly to the same value as the average of the randomly generated structures (after about 15 changes). The maximum value always remains significantly higher than the mean, it drops below PDB quality after only 2 changes. So a high 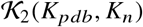 value > 0.75 indicates the structure is likely largely the same as the original structure.

**Figure 7:**
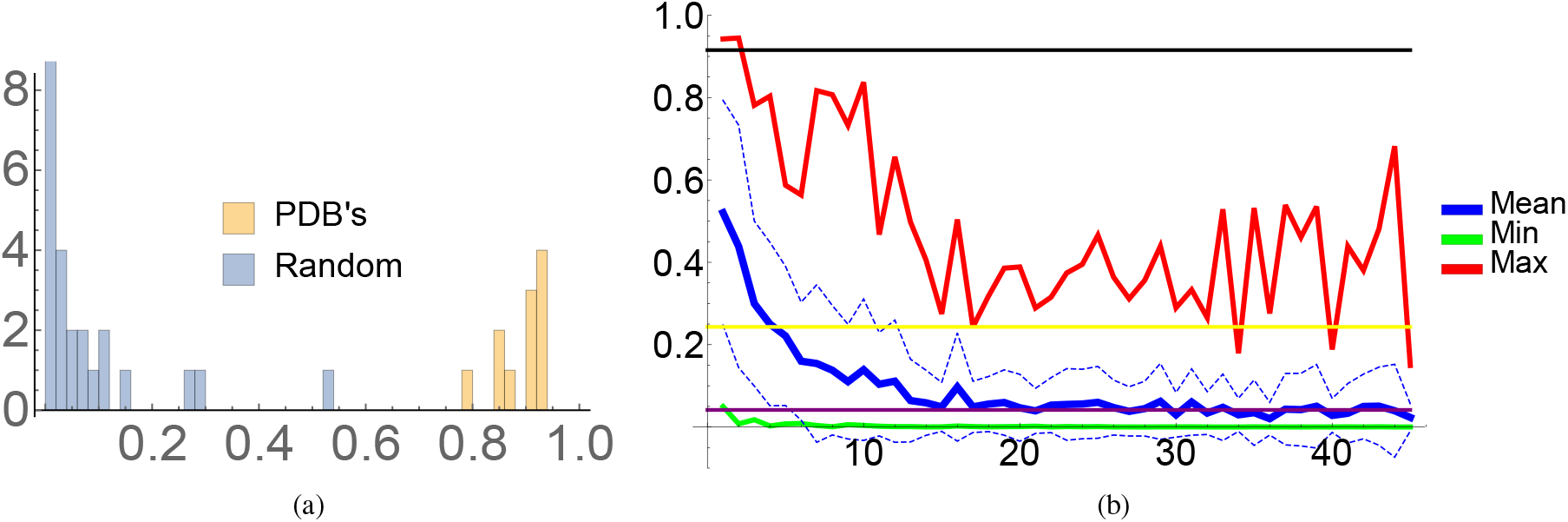
Properties of the (secondary) knot fingerprint statistic 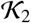 based on variations of the Lysozyme structure. (a) Secondary knot statistics 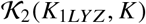 of various structures *K* compared to the curve shown in Figure 6(a). The two distinct sets are Lysozyme PDB’s and random structures with secondary structure alignment to Lysozyme (generated using the CB algorithm). (b) Plots of the mean, maximum and minimum value of the 50 secondary knot statistics comparing the 1LYZ structure and the same structure subjected to *n* random changes in its secondary structure. The dotted lines show 1 standard deviation from the mean. The black line is the average of the PDB structure secondary fingerprint statistics (see (a)) the purple line the Random structure average (crossing the mean at about *n* = 15) and the yellow line the average of secondary fingerprint values for models which fit the experimental data (crossing the mean at about *n* = 3).

### Experimental data fitting

The following chi -square statistic 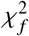 is used to to asses the fit quality of a model predictions

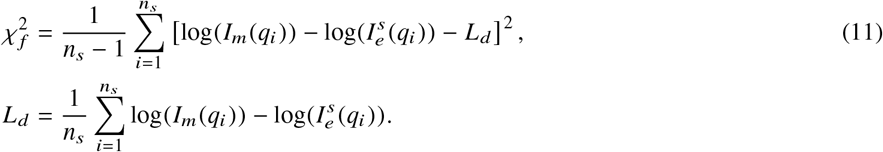

Where *n*_*s*_ is the discrete number of points on the domain *q* ∈ [0, 0.4] on which the scattering is sampled (a commonly used domain *e.g.* (7)). *I*_*m*_ is the model scattering calculated using the Debye formula (5) and 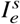, the smoothed experimental data (smoothed using the procedure described in (44)). The factor *L*_*d*_, which will superimpose identical curves which differ by a translation, is used because the protein concentration can only be measured with relatively low accuracy (6, 7) (when taking a logarithm of the data a scaling factor becomes a vertical translation). In addition, to prevent chemically unreasonable conformations, a penalty is applied if the C^*α*^-C^*α*^ distance of ≤ 3.8 occurs for any pair of non-adjacent C^*α*^ positions, this quantity is labelled *χ*_*nl*_ (see Methods and Materials). The initial model is optimized as described above until 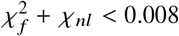. Values below this threshold represent an excellent fit to the scattering data, as shown in Figure 8(d). This value is based on a visual interpretation of fit quality and a comparison to other studies.

**Figure 8:**
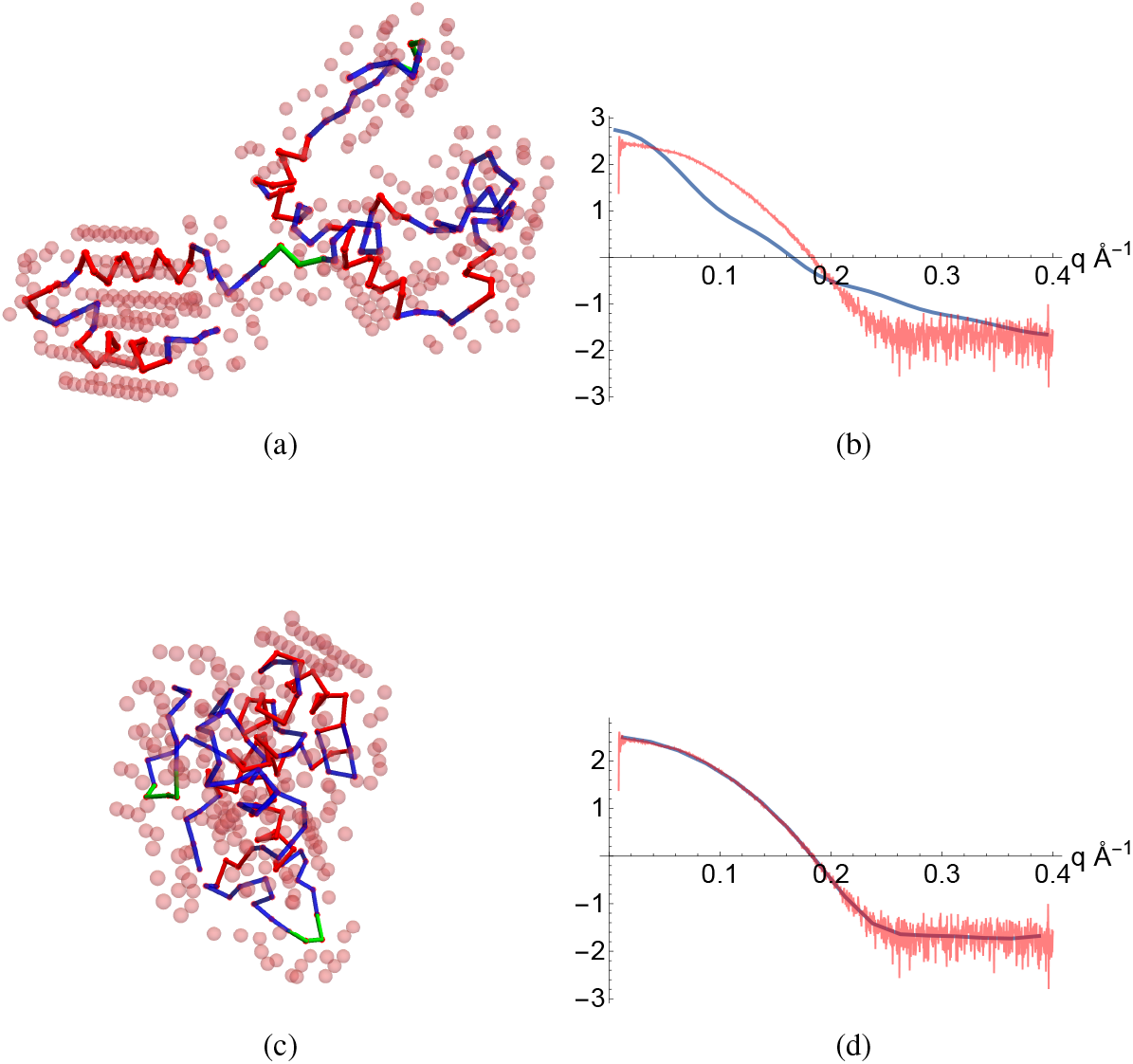
Figures illustrating the fitting process. (a) An initial configuration of the backbone based only on the secondary structure assignment of Lysozyme (PDB:1LYZ). Also shown as spheres are the molecules of the hydration layer. (b) The model scattering curve compared the BioSAXS data. (c) a final structure (and hydration layer) obtained from the fitting process and its model scattering curve now fitting the BioSAXS data well (d).

## RESULTS

In the previous sections a protein model was developed for the interpretation of BioSAXS data which can derive a realistic tertiary fold for the protein using **only** its secondary structure annotation. The process for obtaining a model is summarized in Figure 8. First an initial structure is generated (this is done in various ways in what follows) and surrounded with an explicit hydration layer (Figure 8(a)). A model scattering curve is calculated and compared to the experimental data (b). The curve is then changed by using a Monte-Carlo algorithm to generate new secondary structure units (along with a new hydration shell), thus altering the model’s fold until it attains a sufficiently good fit to the scattering data Figure 8(c) and (d). In what follows we test the efficacy of this process in various stages.

### Validation of the backbone curve and water model

A typical procedure for using small angle scattering data to validate a structure is to use an appropriate set of PDB coordinates as a model which is then fit to the scattering data by adjusting the shape of a hydration layer (via some model of this layer). This is the validation method used in CRYSOL (6) and FOXS (7) ans AQUASAXS (8) for example. A variant of this validation process was performed in section 3 of the supplement. A realistically constrained parameterised hydration layer scattering model was coupled with the explicit hydration layer shell introduced in section to fit an appropriate PDB structures to scattering data. In each case the parameters of the hydration scattering model differed and we proposed to use an averaged scattering model of these individual cases in ab-initio scenarios. If this average scattering model is then re-applied to the PDB structure and explicit hydration shell we do not obtain a sufficiently good fit to the scattering data (although it is not too far off).

The aim of this section is to show that we can use this averaged model and distort an initial PDB model in order obtain a high quality fit to the scattering data whilst still retaining a sufficiently realistic structure (within a few angstroms on average). This demonstrates ab-initio technique proposed here contains within its potential prediction population a high quality representation of the actual protein structure.

To perform this test we selected three pairs of proteins and crystal structure : Lysozyme (PDB:1LYZ), Ribonuclease (PDB:1C0B) and Bovine Serum Albumin (BSA, PDB:3V03, selecting a monomer unit) and scattering data obtained from the SAS database (35). We used the PDB coordinates and secondary structure assignment as an initial input into the algorithm, then we altered each secondary section individually using Monte Carlo sampling of the *κ*-*τ* distributions and the CB algorithm generate new structures. Then, using the hydration layer and scattering model, scattering curves were generated until a suitable fit to the scattering data was obtained.

### Lysozyme and Ribonuclease

Examples of the derived models obtained for Lysozyme are compared to sections of the original PDB in Figure 9, we compare sections for visual clarity. Typically the structures are nearly identical with only the occasional slight deviation in the geometry of some of the linker sections. This similarity is reflected in both the RMSD measures (calculated using the Biopython class (45)) and the knot finger print statistics, as shown in Figure 10(a). As one would expect both indicate excellent fits to the structure. There is a correlation of 0.3 between the two measures, a value on the edge of weak and reasonable (we shall see shortly in the Ribonuclease case a much stronger correlation so we view this as indicative of a relationship between the two measures). The results for Ribonuclease were very similar and the fit statistics are shown in Figure 10(b); again there is also a clear relationship between the knot fingerprint statistic and the RMSD measure, in this case the correlation is very strong, −0.8. We see this the correlation between the two measures as further justification of the knot statistic’s appropriateness as a measure of structure.

**Figure 9:**
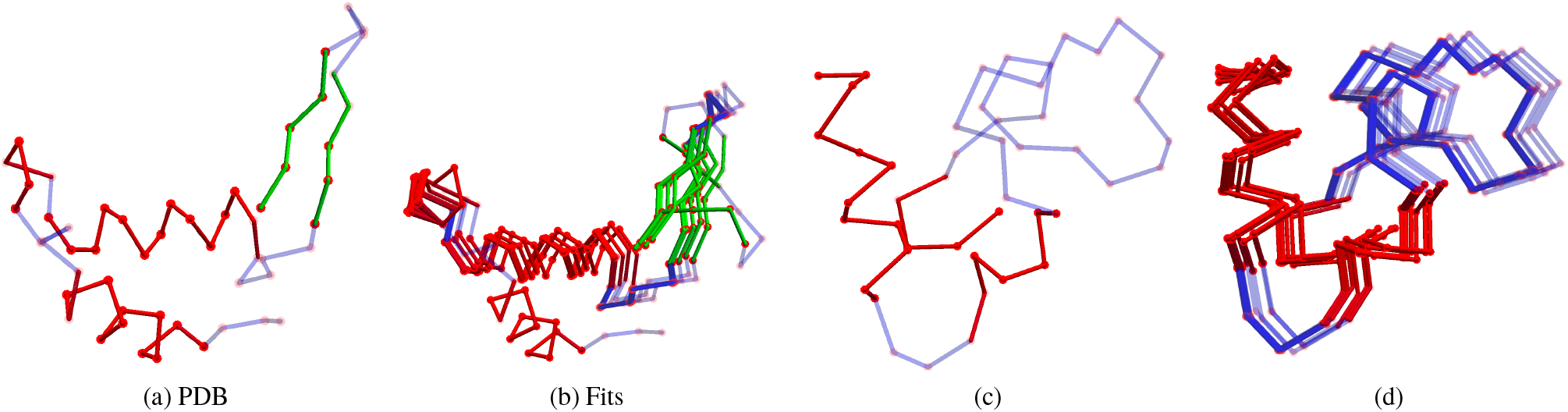
Sections of the 1LYZ PDB structure and example fits obtained by fitting our model to the scattering data. Panels (a) and (c) are subsections of the PDB, (a) has the sheet. Panels (b) and (d) are composite visualizations of the predictions.

**Figure 10:**
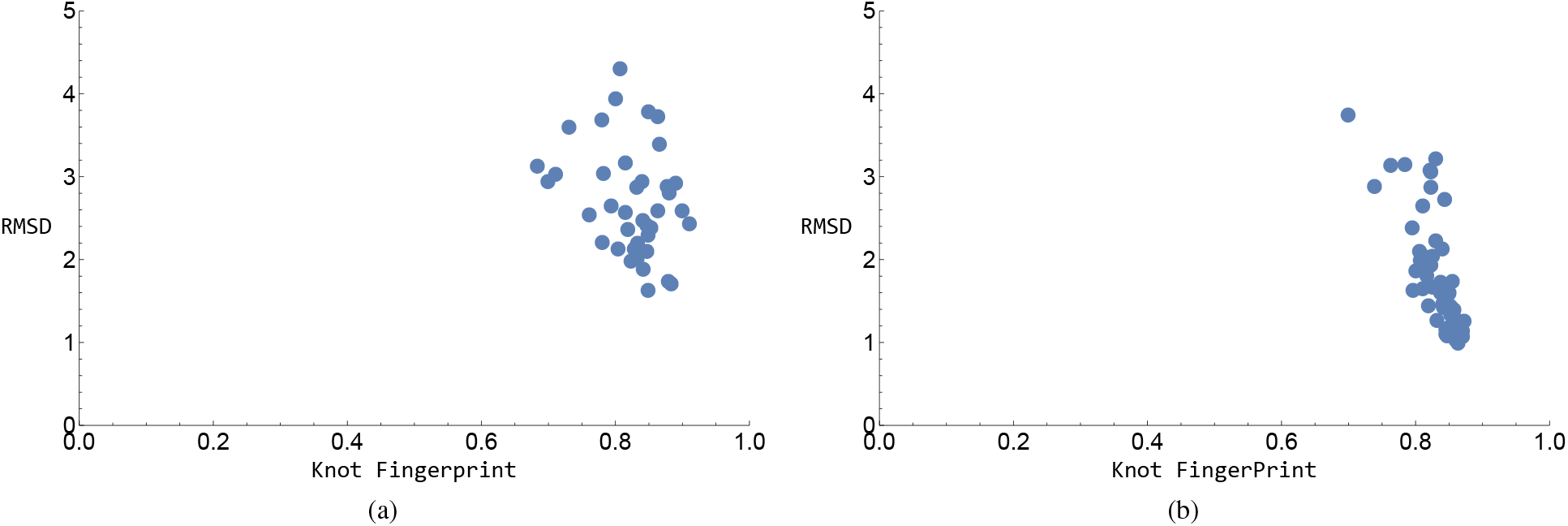
A comparison of RMSD measures and Knot fingerprint statistics 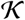 for fittings of the model to scattering data for Lysozyme and Ribonuclease.These results are obtained using the PDB structure as the initial input to the algorithm and are by comparison to that PDB. (a) Lysozyme, (b) Ribonuclease.

### BSA

Example fits to the (parts of the) larger BSA structure are shown in Figure 11, we only display sub-sections as the full molecule is too complex for a clear visual comparison, the sections where chosen at random and are indicative of the general comparison. Once again it is clear the structures are very similar.

**Figure 11:**
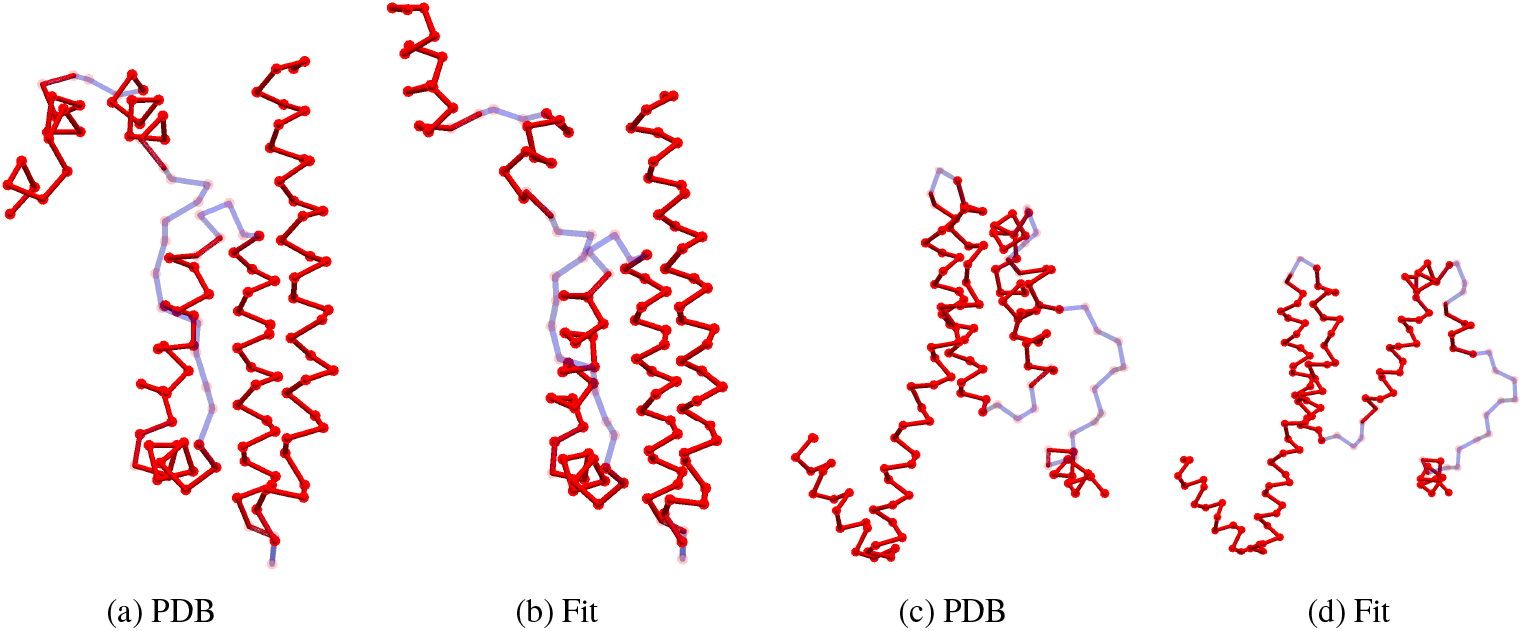
Sections of the 3V03 PDB structure and example fits obtained by fitting our model to the scattering data. Panels (a) and (c) are subsections of the PDB. Panels (b) and (d) are example predictions.

So it is clear the model and method has the potential to correctly predict the tertiary structure of proteins accurately. From a purely ab-initio perspective the question now is how easy is it to get to the correct structure from a random initial guess? This question proves to be more complicated, requiring multiple predictions so for this preliminary study we focus on a single structure, Lysozyme.

### Ab-initio prediction

In the case where no crystal structure is available, the secondary structure prediction based on the sequence alone can be used as a starting point. In order to test this ab-initio method we used the small angle scattering data of Lysozyme to make predictions of its structure. The optimization procedure is similar to the previous section except that an initial conformation of the structure is randomly generated by the CB algorithm. This structure then altered until a suitable fit to the experimental BioSAXS data is found. Once again we use the 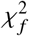 statistic (11), but this time with additional constraint on the potential search space, contact predictions, based on a large number of homologous sequences. Data from the Raptor X web server (46) for the Lysozyme primary sequence were obtained. The C^*α*^ pairs with the 10 highest correlations were selected. An extra potential *χ*_*con*_ was added to the optimization statistic to ensure the distance between these pairs was restricted to be within 5 and 15 Å. If *l* = 1, … *n*_*c*_ labels the *n*_*c*_ pairs of constrained points with mutual distances 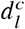 then the quality of contact match *χ*_*con*_ is defined as follows:

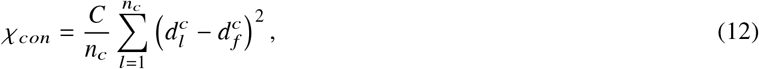

with *C* a constant and 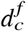 a reference distance (7 was used in this study). The value of *C* controls the likely variation in the distances 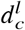, a value of *C* = 0.01 in this study was found to give good results.

The results of the ab-initio fitting procedure are shown in Figure 12(a). The RMSD and knot fingerprint statistics, compared to the 1LYZ crystal structure are shown. The first observation is that the best knot finger print statistics are comparable to the lower end of the from-PDB predictions obtained in the previous section. The second is that these correspond to the best RMSD measures. The apparent correlation between the two measures seems to remain for knot fingerprint statistics above 0.6. However, there is a gap between the best RMSD for the ab-initio predictions and those derived form the PDB structure. This is to be expected as the knot statistic is more tolerant of differences which preserve the entanglement (the general geometry of the fold). This difference can be seen visually in Figure 12 (b) and (c) which respectively represent the first 10 secondary structure sections of the 1LYZ crystal structure and the best fit ab-initio prediction (the one closest to the PDB predictions in Figure 12). The same fold-back of the two significant *α*-helical sections is present in both cases, as is the fold back of the *β*-sheet (although the variability in strand geometry allowed in the algorithm means they aren’t identical). Further the relative orientation of this helical pair and the strand section is present is the same in both cases. So overall the basic fold geometry is correctly predicted which is why the knot statistic is so close to the PDB values. There are, however, a number of sections with some reasonably significant distance differences, for example the linker section joining the two helices; this means a bigger difference in the RMSD measure. For the kind of resolution available in Small angle scattering experiments we argue the knot statistic is a more appropriate measure of the accuracy of the prediction. One can see a similar conclusion can be applied to the rest of the molecule shown in Figure 12(d) and (e) for the PDB and fit respectively.

**Figure 12:**
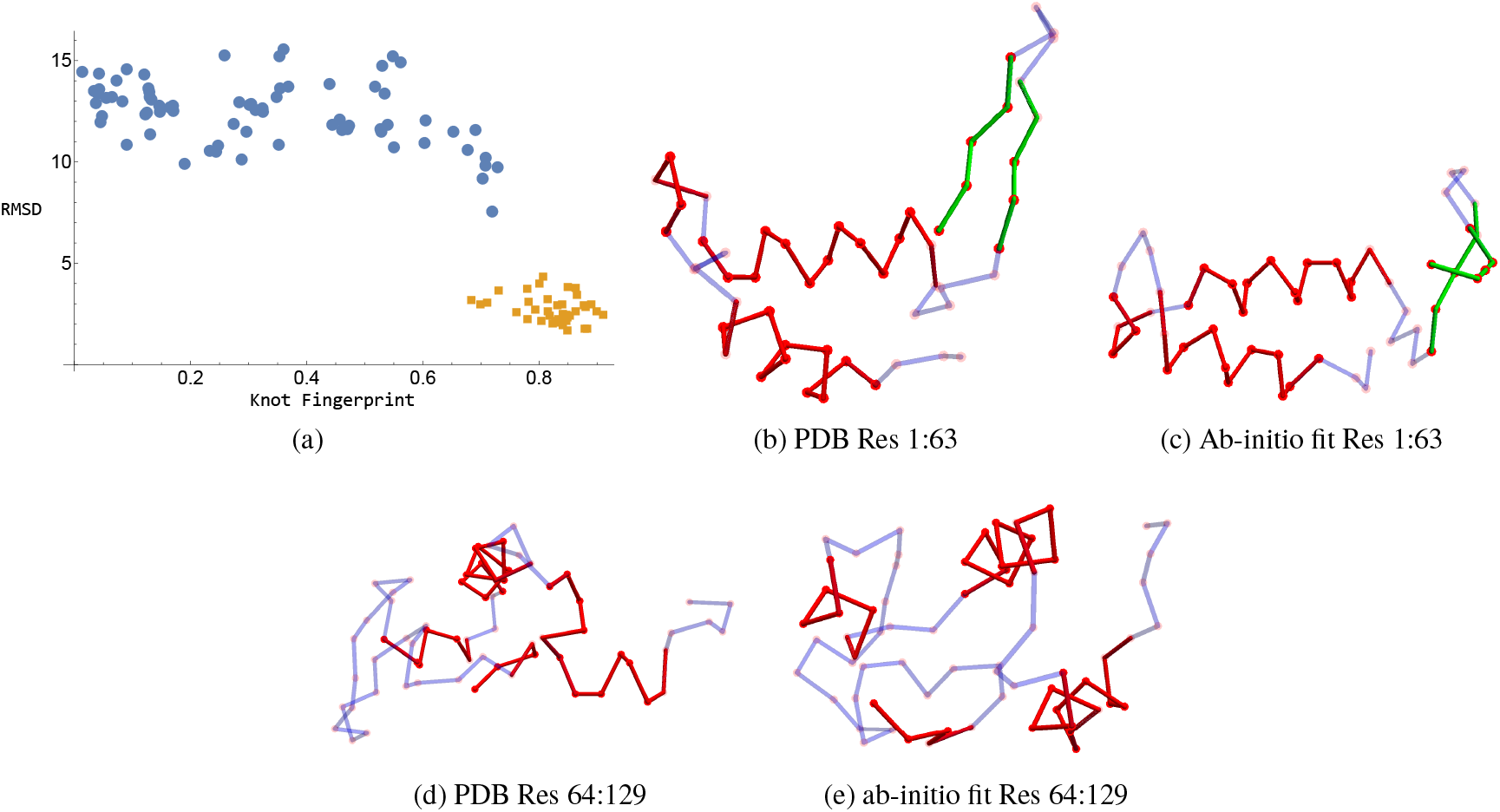
Ab-initio predictions for Lysozyme based on sequence data alone. Panel (a) depicts the RMSD and Knot statistic 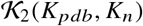 values for the predictions *K*_*n*_, these are indicated as blue circles with the form-PDB data (Figure 10) shown as brown squares for comparison. Panel (b): secondary structure sections 1-10 (residues 1-63) of the 1LYZ crystal structure. Panel (c): secondary structure sections 1-10 of the best ab-initio fit. Panel (d): secondary structure sections 11- of the 1LYZ crystal structure (residues 64-129). Panel (e): secondary structure sections 11- of the best ab-initio fit.

### Objective prediction comparisons

Using only the protein sequence for secondary structure prediction and BioSAXS data we have been able to obtain tertiary structure models which can be observed and quantified to have a significantly similar fold geometry (topology) to the Lysozyme structure. However, a large number of predictions have knot statistics which suggest the structure’s fold topology differs significantly from that of the crystal structure (Figure 12(a)). The target applications for this method will be unknown structures and it must be established whether one could have identified these were “ bad” predictions without the knowledge of the underlying structure.

To differentiate predictions we should seek objective structure comparison measures which do not depend on comparison known structural information (i.e. not to the PDB). One example would be the contact prediction statistic *χ*_*con*_. This is objective in the sense that it only relies on sequence predictions, and would generally be available in target applications. A scatter plot of the knot statistic indicates the high quality ab-initio predictions (high 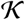) are less likely to have a high *χ*_*con*_ than the worse predictions, see Figure 13(d). If we were to run a significant number of predictions and then say select only those below the mean *χ*_*con*_ value then most of the high *χ*_*con*_ predictions remain, this could be a first means of filtering the predictions, although we see it will still leave “bad” predictions so further analysis is required.

**Figure 13:**
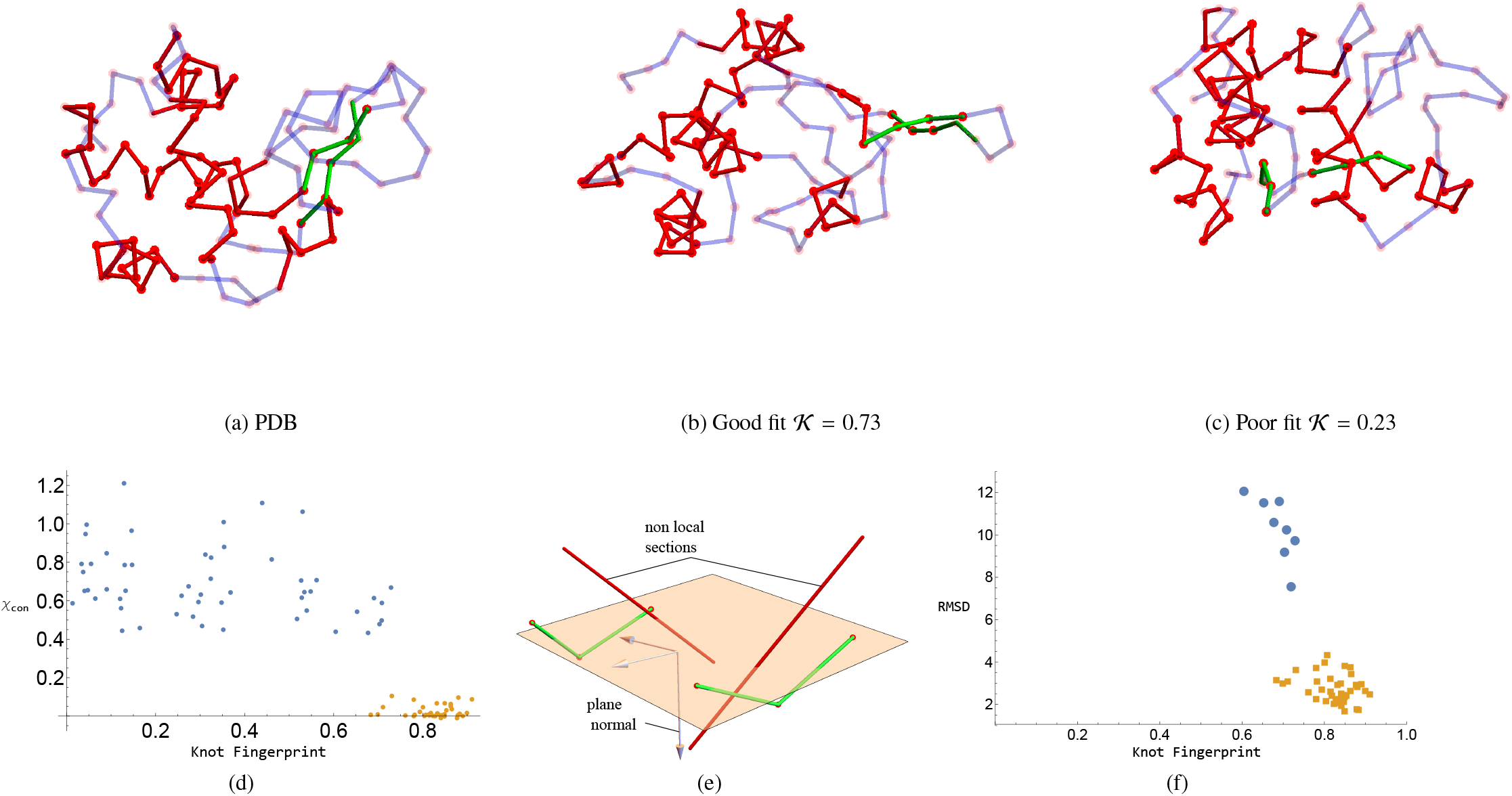
Comparisons of high 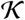 and low 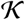 Lysozyme predictions. Panel (a) is the PDB:1LYZ crystal structure. (b) A high quality fit 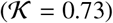, (c) a low quality fit 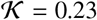. (d) a comparison of the contact constraint *χ*_*con*_ and the knot finger print, the blue points (with larger values) are for the ab-intio fits and he brown dots are the from PDB fits. (e) two (green) sections of a sheet from Lysozyme model. A plane and its normal bi-secting the strand sections is shown, also shown are two sections of the rest of the molecule which bisect the plane between the two strands. (f) the fingerprint-RMSD comparison plot with the screened ab-initio predictions.

### *β* sheet model variations and the power of knot statistics

Figure 13 shows the full 1LYZ crystal structure (a), a high 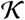 model (0.73) (b) with RMSD 9.21 (by comparison to the crystal structure) and a low 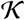 model (0.23) (c) with RMSD= 10.8. So there is a relatively small difference between the two prediction’s RMSD measures, but a significant one as measure by the knot topological method. One clear difference is the isolation of the *β*-sheet. In both (a) and (b) the sheet is at one edge of the structure, whilst is (c) it is closer to the alpha helical structure, and further because its constituent strands of the prediction shown in (c) are not sufficiently closely related there appears to be a section of *α*-helix passing between them. This is a significant difference in entanglement detected by the knot based measure. However an inspection of the structures indicated that the better performing structures (in terms of their fingerprints) tended to have tighter and more isolated *β*-sheets. To try to quantify this we created two mathematical measures. The first measure is the mean distance between sequentially paired C^*α*^ atoms (this sequential dependence can be determined by distance measures and does not need a pre-determined knowledge of the strand orientation). We calculate this value for all predictions and choose those say less than the median value. The second is a discrete test as to whether any other section of the molecule passes “between the sheet”. We approximate a plane for the sheet as indicated in Figure 13(e) and then determine if any other arcs of the main C^*α*^ chain pierce this plane, this this does occur we simply reject the structure as being physically unrealistic (as is the case in Figure 13(e)). Both are objective measures.

When the combination of sheet measures and the contact prediction cut-offs are applied we are left with a significant proportion of the high quality fits, including the one with the lowest RMSD (Figure 13(d)). Crucially all the lower quality fits are filtered out. It should be noted that one of the high quality 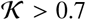 predictions was lost during this filtering process, on the basis that its mean sheet distance was too high. This underlying selection mechanism should be generally applicable being based on basic principles, so there is an indication it will be possible to produce a general post-hoc selection procedure. In future it might be also be useful to use information such as sulphide bonding and hydrophobic exposure to further classify predictions.

### Application to a novel protein with unknown 3D structure: the human SYCE1 core

Based on the success of utilizing contact predictions to constrain potential models we applied the algorithm on the structural core of the human SYCE1 protein, a tethered construct where the sequence is repeated to allow formation of an extended anti-parallel coiled-coils with two short additional helices at each end that could fold back to form a small 3-helix bundle. The secondary structure of the tethered protein construct resulted in eight stretches of alpha-helices where based on the heptad repeats helices 2, 3, and 4, can be aligned to helices 6, 7, and 8 corresponding to the same sequence, respectively in an anti-parallel fashion. This resulted in 14 close contact predictions between helices 2 and 8, and helices 4 and 6, respectively, as shown in Figure 14.

**Figure 14:**
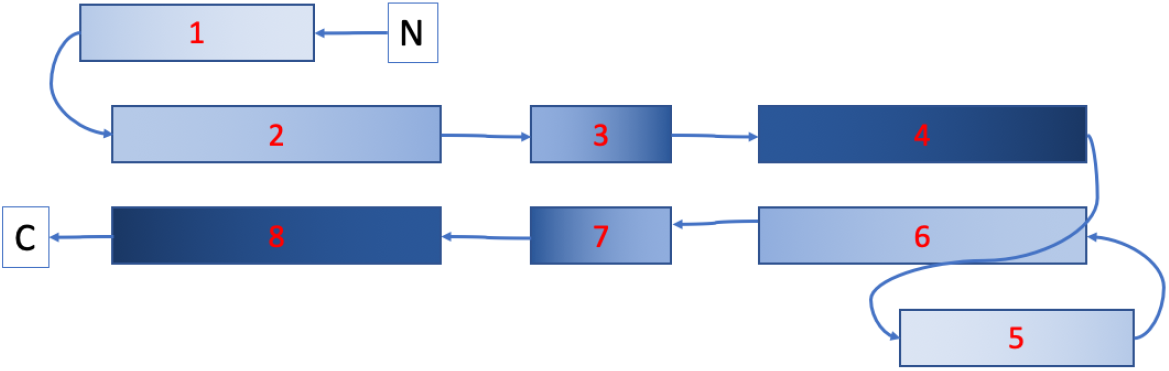
Schematic drawing of the SYCE1 construct with each box corresponding to one predicted alpha helix. The SYCE sequence of approximately 120 amino-acids corresponding to helices 1-4 is duplicated and linked by a tether to a repeat of the same sequence comprising of helices 5-8.

### Deriving the models

Based on the sequence and secondary structure predictions (a combination of those of Raptor X (46) and HHPRED (47)) 40 initial configurations were generated using the CB algorithm. An example is shown in Figure 15(a) along with its hydration layer, its scattering curve is compared to the experimental data (from (24)) in Figure 15(b). As shown the fitting is limited to the domain *q* ∈ [0, 0.3]^−1^, which balances the twin consideration of a sufficient resolution and reliable signal to noise ratio. Using monte-carlo optimization the structure is altered until a reliable fit 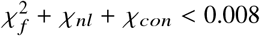 is obtained, where the potential *χ*_*con*_ is based on the contact predictions described above. One such model is shown in Figure 15(c) along with its scattering curve in Figure 15(d). The identical chains of the structure have folded to lie (nearly) parallel with the end termini occupying a local neighbourhood. Two example models for which 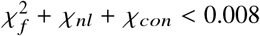 are shown in Figure 16(a)-(b). Figures 16(c) and (d) indicate one of the coiled coil structures and depict the pairwise distances associated with the contact prediction terms *χ*_*con*_. All models share the elongated bend shape with a anti-parallel coiled-coil arrangement of helix 2-4 to 6-8, respectively. The first helix in each helix (helices 1 and 5, respectively) show different orientations which reflect the expected conformational flexibility of the protein in solution. Importantly, the central coiled coil (made of helices 3 and 7, respectively) is not based on the constraints given a-priori but is entirely based on the optimization against the experimental data. Although a bead model results in a similar overall shape (24) our methods is able to derive a more detailed molecular model with distinct structural features such as a coiled coil.

**Figure 15:**
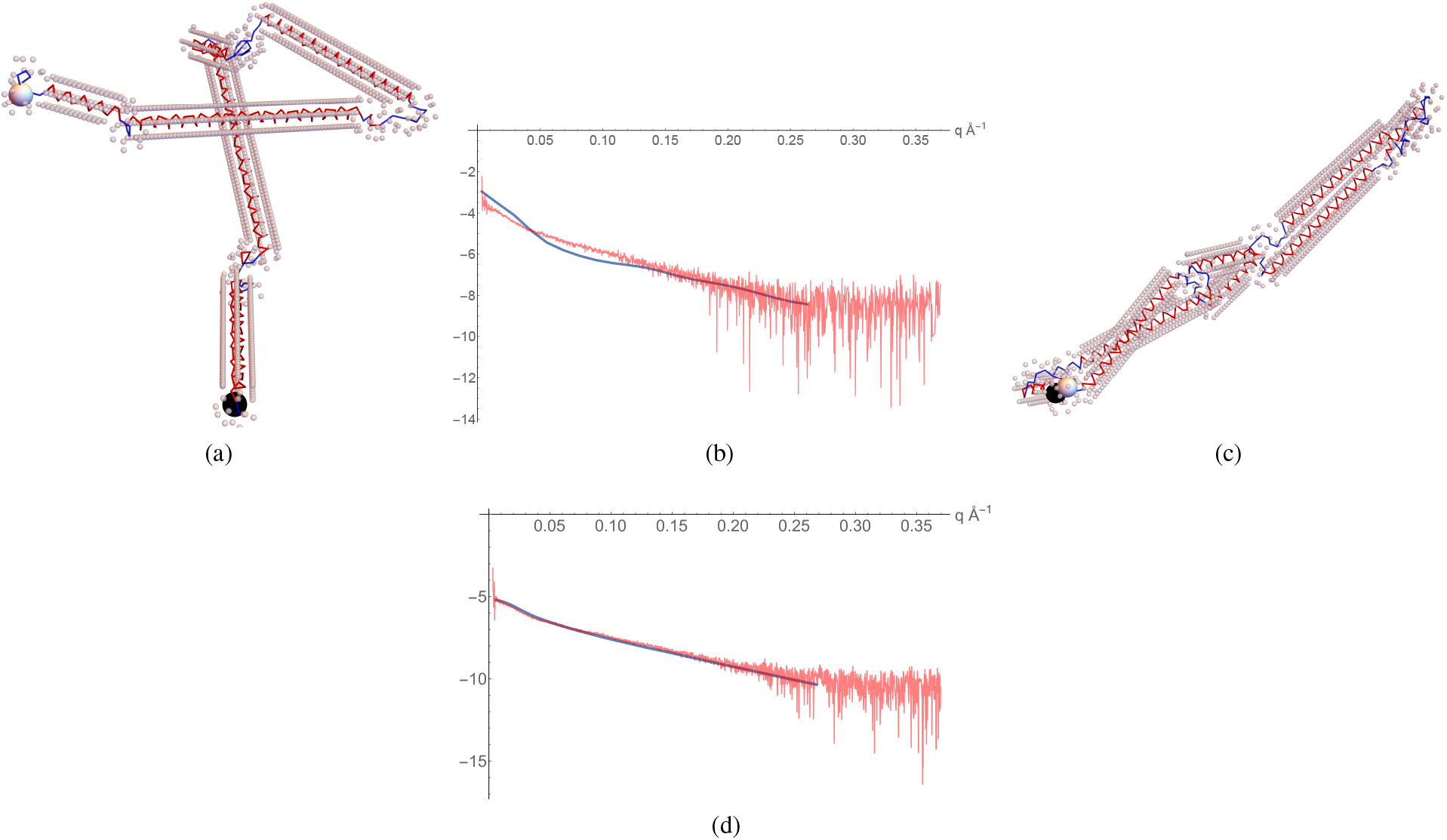
Illustrations of the optimization process used to obtain the model predictions forthe structural core of human SYCE1. Panel (a) An initial configuration of the backbone based only on the sequence data shown in Figure 14. Also shown as spheres are the molecules of the hydration layer. Large black and white spheres indicate the end termini. (b) the scattering curve of the initial configuration (blue) over-layed on the scattering data (red). (c) the model prediction for which 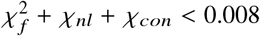, the end termini are next to each other. (d) the final scattering curve compared to the experimental data.

**Figure 16:**
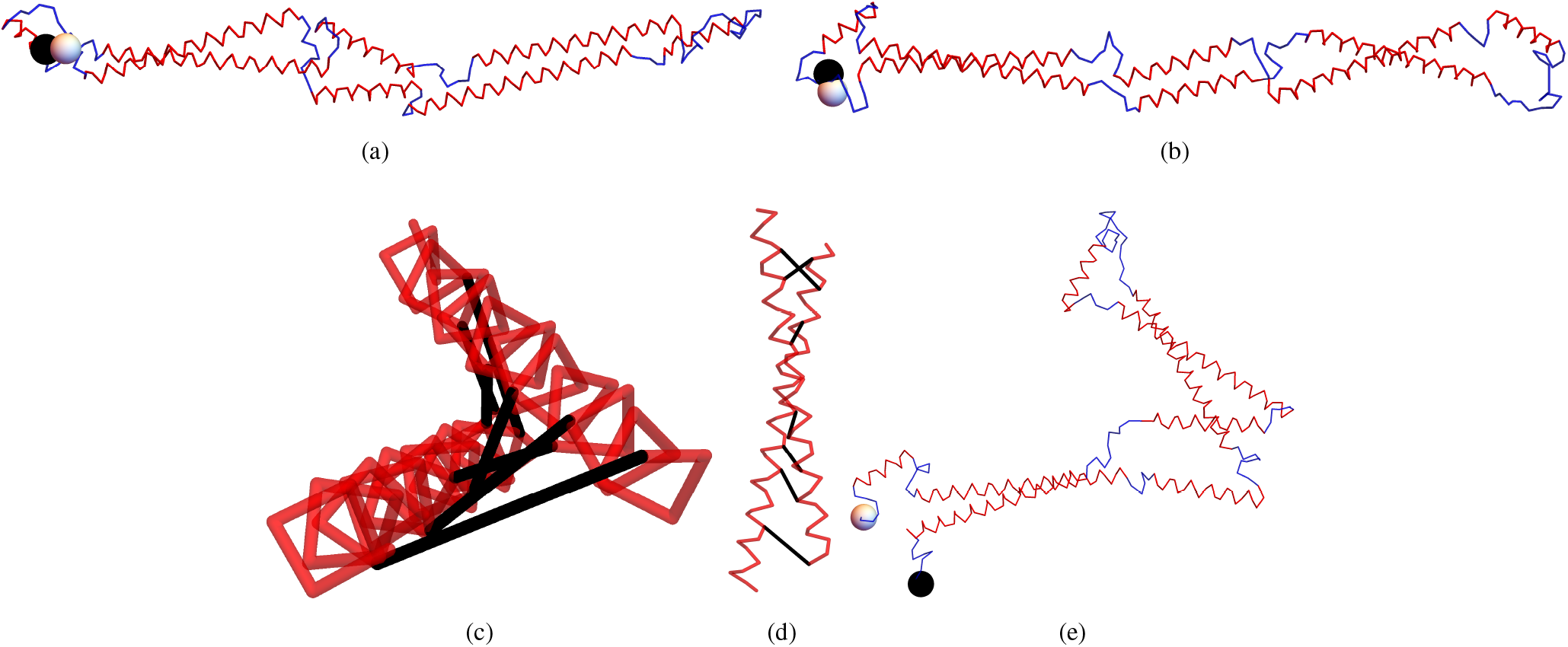
Illustrations of the model predictions. (a)-(b) are all model predictions. (c) One of the coiled coil units of (a) with black tubes representing the contact prediction distances, as see along the axis of the unit. (d) the tilted helical structure of the coiled coil unit. (e) a model obtained by minimising the chi-squared measure *χ*_*nl*_ + *χ*_*con*_ only.

### The experimental scattering data is crucial to the prediction quality

One might ask if the contact predictions alone were sufficient to predict the structure, since they are crucial to forming (some of) the coild-coil structure. To test this we derived models by minimising the chi-squared measure *χ*_*nl*_ = *χ*_*con*_ (*i.e.* ignoring the scattering data), a typical example is shown in Figure 16(e) The outer *α*-helicies are present as the contact prediction constraint *χ*_*con*_ force these structures to form. However the whole structure is significantly folded. This folding was found to be a typical property of models obtained by minimsing only *χ*_*nl*_ + *χ*_*con*_ and the degree of folding was far from consistent. The clear effect of further enforcing the model fit the scattering data is two-fold, first straightening out the whole structure and secondly, in doing so, developing a coiled-coil geometry in the middle of the structure.

### Fitting to the scattering data and contact predictions is not straightforward

As a final note we note that of the 40 initial structures generated, only 5 obtained a suitably low combined chi -squared statistic (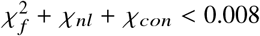). All 5 structures, two of which are shown in Figure 16, were basically identical in this case (comparative ‖ values > 0.9) so there was not need for any post-hoc structural comparison analysis. By comparison all 40 lead to models for which *χ*_*nl*_ + *χ*_*con*_ < 0.008. As we have just seen there is significant value in the extra information provided by the scattering data. The difficulty with obtaining suitable fits indicates that in the future more advanced optimization techniques than a straightforward monte-carlo search may be needed.

## DISCUSSION

This paper details in depth the development of a tertiary structure model for ab-initio model generation from BioSAXS data. A number of key points have been demonstrated with regards to its potential use to the structural biology community. Firstly, if the method takes as input a structure significantly similar to the target structure *e.g* a homology model or an incomplete (core) model for a structure or a model from homology, then it will likely find a highly accurate fit to the structure. Secondly given a near complete absence of tertiary structural information, save that available from sequence data, the technique can generate realistic representations of the structure’s fold. Further, in this ab-initio scenario there is the potential to reliably separate realistic predictions from those which are no biologically plausible, by both constraining the fitting procedure and applying post-hoc filtering measures.

With regards to comparisons to existing techniques there are two categories to be discussed. The first is the set of different experimental used to derive structures in the protein data bank. The predictions from our methods, applied to small angle scattering data, can be near this level of quality **if** a reliable initial structural prediction is provided, this was demonstrated in section where we say structures with RMSD measures of the order of accuracy of a of a typical structure on the Protein Data Bank for *α* carbon positions. In a purely ab-initio scenario our results in section indicate it is currently difficult to obtain this level of accuracy on a a reliable basis. However, as indicated in Figure 13(d), there is some indication that, if extra constrains such as contact predictions from homologous sequences can be enforced to a high degree of accuracy, that there is the potential to reach similar levels of structural resolution to these alternative experimental techniques.

The second comparison would be to SAXS specific ab-initio techniques for interpreting BioSAXS data. These include the bead based models such as GASBOR and DAMMIN/DAMMIF (13, 48). It is not possible to make a direct comparison between the methods as we did for the crystallographic predictions as the nature of the prediction is different. Neither method makes explicit predictions of the tertiary structure of the molecule as being constructed form secondary structural elements. Both are composed of effective scattering beads, the DAMMIN model aims to predict the volume occupied by the molecule by creating a cloud of beads whilst GASBOR does aim for a structure with a chain like nature constraining bead-bead distances, but there is no explicit secondary structure in the model. In the case of Lysozyme on can verify that the PDB structure cane be superimposed onto(into) the GASBOR and DAMMIN predictions (see *e.g.* (13)), one can see that our predictions also have this property of occupying a similar volume to the crystal structure in Figure 13(a)-(c). The advantage of our model is that it also makes an explicit prediction for the fold geometry of the secondary structure elements.

The Atsas package does allow for the interpretation of bead models with tertiary structure through the use of the CORAL package (48). Given known structures the package attempts to fit the structure into the bead model with a mixture of known (manually assigned) and unknown elements. This procedure was performed in (24) provide evidence that the SYC1E core modelled in section was a coiled-coil domain. Two coiled-coils were superimposed on a bead model with CORAL providing an additional linker section to join them. Our model simply uses the sequence data to determine the secondary structural elements, then it is able to try millions of differing (physically realistic) folds which and tests **each time** if they satisfy the scattering data, a much more direct and exhaustive test, which relies on far less user input. What is interesting is that this technique predicts an additional coiled-coil domain at he structure’s centre, owing to the sequence interpretation splitting of the helical units. We feel our method is, in the long term farm more flexible and amenable to automated structural evaluation with its main comparative advantage is the potentially exhaustive automated search of the potential search space.

## CONCLUSION

As a solution-based technique, BioSAXS can provide structural information for targets where crystallisation proves difficult and can also allow data collection in a more natural environment than techniques such as crystallography and cryo-EM. Additionally, SAXS is not limited by protein size, as is the case for cryo-EM and NMR. So there are clear advantages of being able to develop the techniques for interpretation of this data in an ab-initio setting which improve on the levels of structural detail provided by the bead models currently popular.

In this paper we have shown that curve representation with hydration shell provides a molecular model for BioSAXS data with fits as good or better than traditional bead and envelope models. Unlike these models our model includes a complete secondary and tertiary model description. Importantly, starting from random models that only take secondary structure information and sequence-dependent distance constraints into account, a physically meaningful 3D model can be obtained by fitting models against the experimental data. That this is possible is due to the fact that the model is described with far fewer parameters compared to even a coarse-grain model that required three coordinates for each amino-acid **combined** with use of geometric constraints for regular secondary structural elements.

In order to show the potential of this ab-initio technique it was applied to a tethered core component of the human SYCE1 protein, for which no high-resolution structural data is available. The model derived was based on sequence information **alone** match those of a model that was previously reported in (24). where the model was based on manual inspection of the sequences coupled with the fitting of ideal coiled coil segments to experimental scattering data. Importantly, whilst the previously modelled structure includes two coiled coil segments, the model derived here recognised that this was the minimum number of segments required to explain the curved structure and that the true structure could consist of multiple coiled coils interrupted by short linkers. Thus, our novel ab-initio method has successfully generated a highly plausible model from experimental scattering data without the need for any more than minimal manual evaluation. This facility will be crucial for ab-initio structural determination (from biosaxs data) of larger molecules where it would not be practical to generate structures manually.

Further experimental information such as distance information from any other source can easily been added in the form of additional restraints into the optimization algorithm. The model’s explicit description of realistic secondary structure means additional information, like contact predictions, radius of gyration, hydrophobicity of the chain and disulfide bonding can be employed as model constraints in the future. This will further enhance the accuracy of all potential models, and in particular help the end-user to distinguish mathematically correct but physically less likely models from correct solution. The secondary knot fingerprint statistic developed shows significant potential to evaluate structural similarity of models and hence to further automate this vital validation step.

The two future next steps are (i) the application of this method to multimeric structures where each known monomer structure can initially be treated as rigid-body and then refined in order to account for local changes in solution (ii) the application to larger, de-novo structures where the exact 3D structure remains elusive. The application to homo-multimers is straightforward and requires only minor addition to the existing code, we expect this to be the major initial application of our methods. Due to the limited information content of small-angle X-ray scattering data the ab-initio fold determination will depend on the accuracy of secondary structure prediction combined with appropriately weighted distance constraints such as those discussed above.

## Supporting information

Supplementary model details referenced in the main text

## AUTHOR CONTRIBUTIONS

Chris Prior: Co-conceived the project and wrote the paper. Constructed the model, tested and calibrated it, additionally he developed the knot fingerprint evaluation routine. Owen Davies: Provided experimental data. Provided biological background, reviewed the paper. Daniel Bruce: Assisted with various aspects of the data gathering and processing used to calibrate the model Ehmke Pohl: Co-conceived the project and wrote the paper, provided substantial guidance and biological/biochemical insight towards the model’s construction as well as obtaining the experimental data.

## ACKNOWLEDGMENTS

The authors would like to thank Christina Law for testing the scattering and hydration models. Financial support from the Biophysical Sciences Institute and the Addison-Wheeler fellowship for CP are gratefully acknowledged. We would also like to thank Dr Mark Miller whose knot identification code was used to calculate knot fingerprints as well as Prof Alain Goriely and Dr Andrew Hausrath for their input and advice in the development of the backbone model. Finally we are grateful to Dr Beth Bromley for her help with the contact predictions within the coiled coil of SYCE1.

## SUPPLEMENTARY MATERIAL

An online supplement to this article can be found by visiting BJ Online at http://www.biophysj.org.

