## Supplementary model details referenced in the main text for "Obtaining tertiary protein structures by the ab-initio interpretation of small angle X-ray scattering data"

Manuscript submitted to **Biophysical** *Journal*
Article

### Supplementary material for: Obtaining tertiary protein structures by the ab-initio interpretation of small angle X-ray scattering data

Christopher Prior[1,*], Owen R Davies[2], Daniel Bruce[3], and Ehmke Pohl[3,4*]

[1]Department of Mathematical Sciences, Durham University, Durham DH1 3LE, United Kingdom
[2]Institute for Cell and Molecular Bioscience, Medical School, University of Newcastle, Newcastle upon Tyne, NE2 4HH
[3]Department of Biosciences, Durham University, Durham DH1 3LE, United Kingdom
[4]Department of Chemistry, Durham University, Durham DH1 3LE, United Kingdom

#### 1 RESTRAINING AND GENERATING THE POLYPEPTIDE CHAIN

##### 1.1 Restraining the values of $\kappa, \tau$ and $R$

In practice the geometry of polypeptide chains are restricted according to the restraints on the dihedral angles along the main chain discovered by Ramachandran (1–3). In the following section it is shown that the stereo-chemical restraints of protein structures manifest themselves in a similar way on the $\kappa$-$\tau$ space that is used to describe the $C^{\alpha}$backbone in this model.

To constrain the permissible $\kappa$-$\tau$ values 60 protein structures for which high resolution crystal structures were available were chosen. Using the secondary structure assignments of this set of model proteins the polypeptide chain was split into those designated $\alpha$-helices, $\beta$-strands and the rest not specifically identified were classed as “linkers”. The $\kappa$ and $\tau$ formulae were then applied to all quadruplets of neighboring $C^{\alpha}$atoms. The data are plotted in $\kappa$-$\tau$ space in (a)-(c). Figure 1(a) shows $(\kappa, \tau)$ pairs for all quadruplets identified as belonging to $\alpha$-helices; the vast majority of subsections take values within a small range $\kappa \in [0.4, 0.5]$ $\tau \in [0.18, 0.38]$, $> 97\%$ of the sections fall within this range.

For $\beta$-strands (Figure1(b)) there are three distinct areas where the $(\kappa, \tau)$ values congregate, one in the same region as the $\alpha$-helical structures and two with higher (magnitude) $\tau$ and lower $\kappa$ values, in what follows they are referred to as the left $\tau$ and right $\tau$ regions based on the fact they have positive and negative $\tau$ respectively. There is a similar set of high density regions for the linker sections (Figure 1(c)), though there is also a greater spread of values. In Figure 2(a) (positive $\tau$) and Figure 2(b) (negative $\tau$) typical quadruplets from the left right $\tau$ poolings, are compared to a typical $\alpha$-helix structures, they coil in a tighter and more elongated fashion.

##### 1.2 Along-chain distances

In order to determine reasonable target distances between adjacent $C^{\alpha}$atoms, all distances for the same set of high-resolution crystal structures were calculated and plotted for regular secondary structural elements. The $C^{\alpha}$-$C^{\alpha}$distance $R$ varies very little from a mean value of approximately 3.8Åfor all section types (Figure 3(a)-(c)). This value is assumed to be a fixed quantity in the model.

##### 1.3 Relation to the favored regions of the Ramachandran plots

The distinct clusterings in $(\kappa, \tau)$ space can be shown to correspond to the preferred regions of Ramachandran space. The Ramachandran space is a two-dimensional plot of the “torsion" angles $\phi$ and $\psi$ corresponding to the orientations of the $C^{\alpha}$-N and $C^{\alpha}$-C bonds in each amino acids (note: torsion here is not the same as the $\tau$ we have defined and the angles are not related $\phi$ and $\psi$ used in the curve reconstruction detailed in Methods and Materials). Ramachandran and colleagues (1) showed, using hard sphere approximations, that these values are restrained to occur in pairs which lie in restricted regions of the angle parameters space ($\phi \in [-180, 180]$, $\psi \in [-180, 180]$). This is depicted in Figure (4)(a), which is a scatter density plot taken from Lovell *et al* (4) (note the plot is periodic in both directions). There are three distinct regions of high density which are outlined. Helical sections based on these angle sets were constructed, with dihedral pairs taken from each preferred region, see

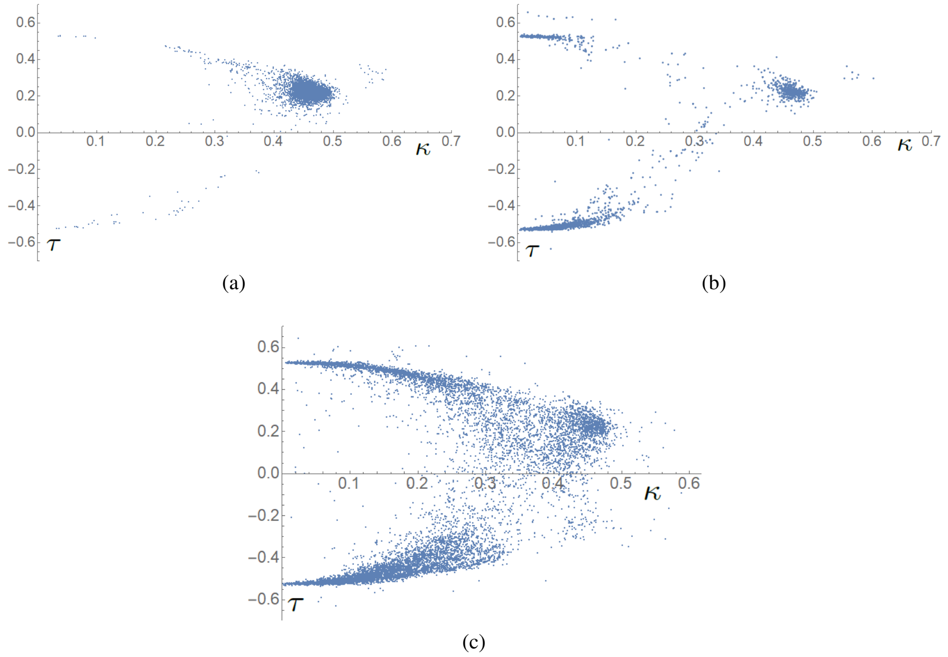


Figure 1: Distributions of $\kappa$ and $\tau$ from a representative selection of high-resolution crystal structures from the PDB. (a) for $C^{\alpha}$'s belonging to alpha helices. (b) for $C^{\alpha}$'s belonging to strands. (c) for $C^{\alpha}$'s belonging to linker sections.

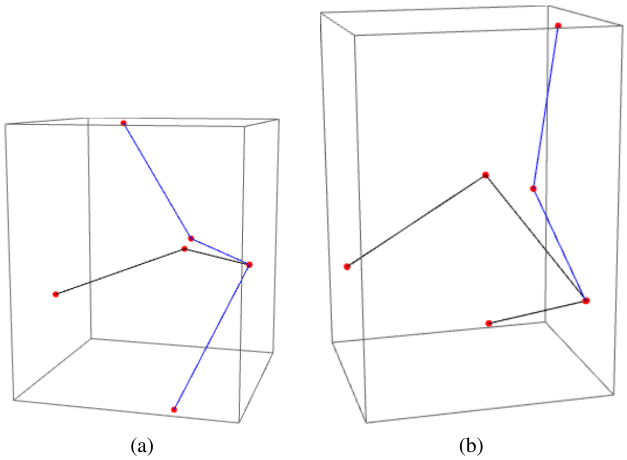


Figure 2: Geometries of strand sections. (a) A positive $\tau$ strand (blue) overlayed on an $\alpha$-helical section (black). (b) a comparison of a negtive $\tau$ (left handed) strand section (blue), again compared to an $\alpha$-helix (balck).

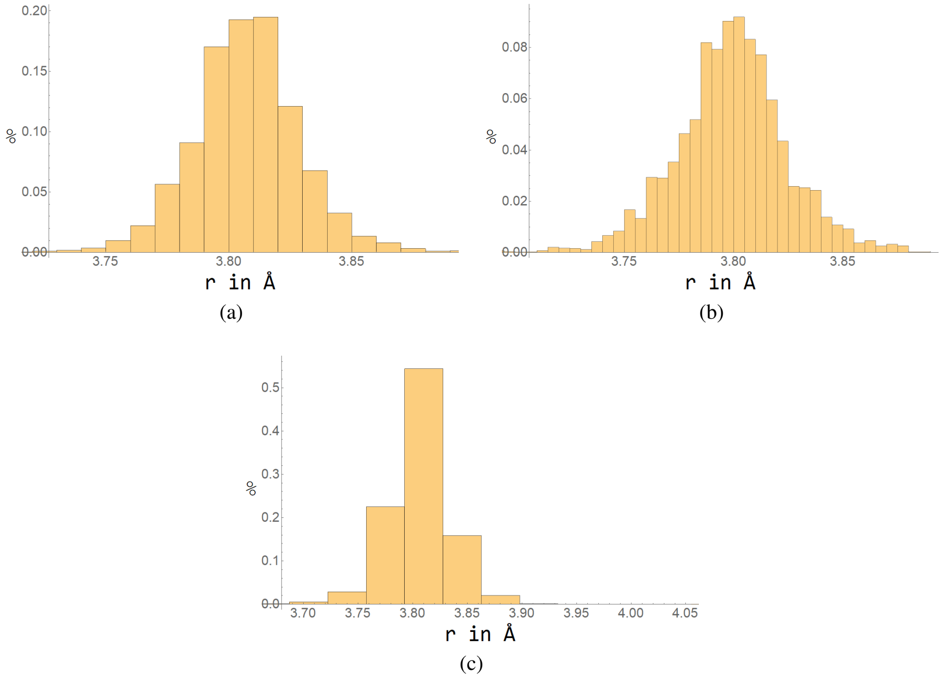


Figure 3: Distributions of $r$, the $C^{\alpha}$-$C^{\alpha}$distance, from a selection of high resolution crystal structures. (a) for $C^{\alpha}$'s belonging to alpha helices. (b) for $C^{\alpha}$'s belonging to strands. (c) for $C^{\alpha}$'s belonging to linker sections.

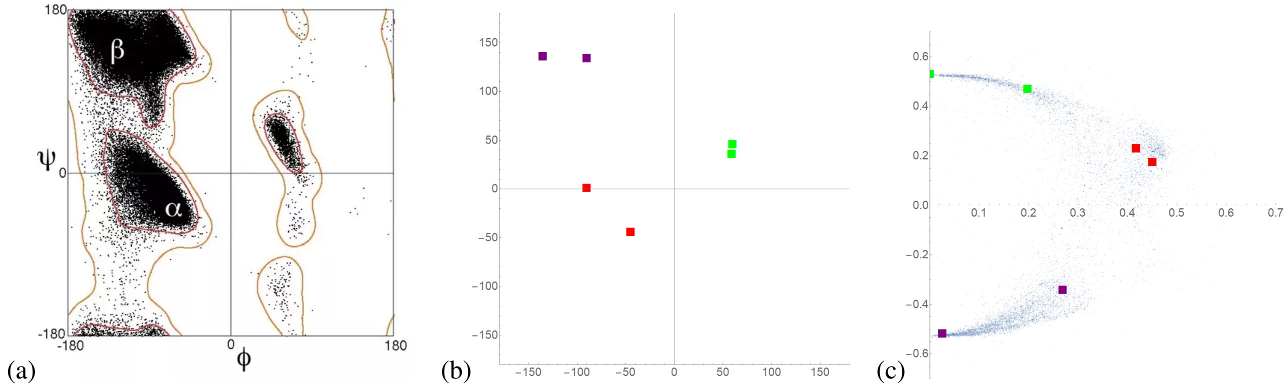


Figure 4: The mapping of Ramachandran space to $\kappa$-$\tau$ space. (a) a general Ramachandran plot the dark regions are density peaks indicating the favoured regions. (b) Dihedral angles used in to create helical curves in order to map to $\kappa$-$\tau$ space. The values are marked as squares. (c) The $(\kappa, \tau)$ values of the helical curves produced using the dihedral angles shown in panel (b).

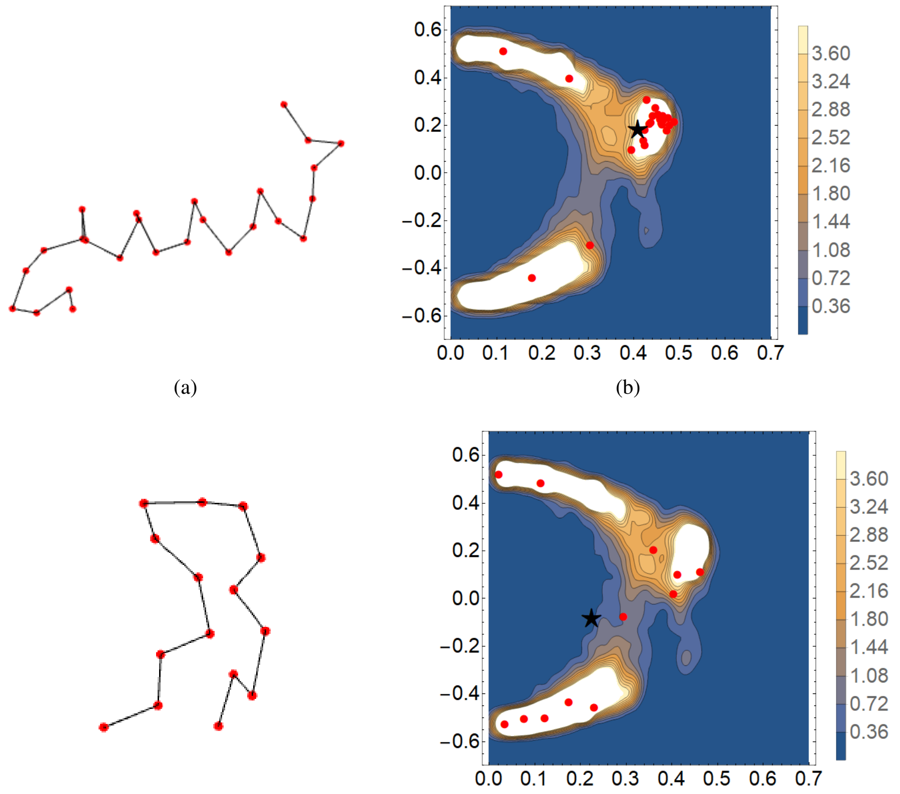


(a) (b) (c) (d)

Figure 5: Example backbone sections and their $\kappa$-$\tau$ values, shown superimposed on a contour plot of a smoothed probability density function of the linker data shown in Figure 1(c) (whose construction is detailed below). (a) is a linker section from the PDB 1OAD (Glucose Isomerase), its interior (the middle points) have a clear helical structure. (b) most of the $\kappa$-$\tau$ values (red points) fall within the $\alpha$ preferred region, as does the average $(\kappa, \tau)$ value (indicated by a star). (c) is also a linker section from the PDB 1OAD, it has a notably non-linear loop structure and its $\kappa$-$\tau$ values, shown in (d), are more evenly distributed amongst the general distribution. Its mean does not fall within one of the three preferred regions.

Figure 4(b). The method for converting dihedral angles into a helical curve is based on some unpublished work by Shah, Tabor, Goriely and Hausrath and can be made available upon request. The $(\kappa, \tau)$ values calculated in each case are shown in (4)(c) superimposed on the $(\kappa, \tau)$ distributions of the linker sections. Each preferred domain of dihedral space can be seen to match to the distinct poolings of $(\kappa, \tau)$ space. So ensuring sections of the model backbone curve are distributed according to the empirically determined $\kappa$-$\tau$ distributions will mimic imposition of the Ramachandran constraints.

#### 1.4 Random coil sections

Often there will be sections of a protein which do not represent regular secondary structure but random coils, these include various turns and loops and often N− and C− terminal residues. In terms of the empirical $\kappa$-$\tau$ spaces this manifests as a section whose $\kappa$-$\tau$ values are almost all confined to **one** of the preferred regions. An example mainly confined to the $\alpha$-helix region is shown in Figure 5(a) (from the PDB 1OAD, Glucose Isomerase (5)), it has a clear helical structure in its interior, its composite $(\kappa, \tau)$ pairs are shown in Figure 5(b), along with their average which indicated by a black star. The fact that the average $(\kappa, \tau)$ point is in the same preferred region as most of the sections individual $(\kappa, \tau)$ pairs demonstrates that the structure is stereochemically constrained. By comparison in Figure 5(c) is section (from the same molecule) whose average $(\kappa, \tau)$ pair is outside of the preferred regions (Figure 5(d)), its structure is far more irregular than the one shown in Figure 5(a).

Random coil sections are treated separately in this model as they have significantly different characteristics to less constrained sections. In a model fitting procedure it would make sense to include a probability that some sections, marked as linker sections are treated as random coils. To include this possibility the mean $(\bar{\kappa}, \bar{\tau})$ values of each section of the linker data were calculated. If this average fell outside of the preferred regions the section was classed it as “mixed", if not it was classed as “linear". Any

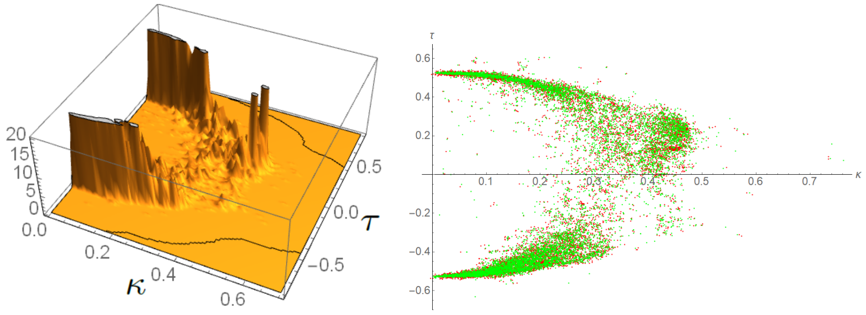


(a) (b)

Figure 6: The probability density distribution of mixed-type linker $(\kappa, \tau)$ pairs. (a) the derived probability density distribution for the mixed Linker data, it has thin, high peaks where the main poolings are. (b) a comparison of the original data and a set of randomly generated $(\kappa, \tau)$ pairs obtained using the PDF shown in (a).

linear sections were then split into three categories based on which preferred region their average $(\bar{\kappa}, \bar{\tau})$ value was contained. Each type occurred with the percentages listed in Table 1 for linker sections. Similarly for strand sections it was found that each type occurred with the percentages indicated in Table 2. The linker sections are much more likely to be mixed than their strand counterparts.

| Section geometry | Percentage abundance |
|---|---|
| Mixed | 87.892 |
| Alpha | 3.823 |
| Left $\tau$ | 6.031 |
| Right $\tau$ | 2.254 |

Table 1: Percentage abundance of geometry types in linker sections.

| Section geometry | Percentage abundance |
|---|---|
| Mixed | 23.536 |
| Alpha | 27.050 |
| Left $\tau$ | 44.985 |
| Right $\tau$ | 4.429 |

Table 2: Percentage abundance of geometry types in strand sections.

#### 1.5 Generating probability density functions for $\kappa$-$\tau$ space.

In order to use the $(\kappa, \tau)$ data to generate potential polypeptide chains probability distributions for $(\kappa, \tau)$ were generated for each structure type. As the underlying probability function is unknown, a kernel smoothing technique was used (see *e.g* (6)). The basic idea is that there is some function $f(\kappa, \tau)$ which gives the probability density function (P.D.F) of obtaining pair $(\kappa, \tau)$. An estimate $\hat{f}$ is generated from a set of data $(\kappa_i, \tau_i)$, $i \in 1, \dots N$, where $N$ is the number of data points, using the following smoothing kernel formula

$$\hat{f}(\kappa, \tau) = \frac{1}{Nh} \sum_{i=1}^{N} D\left(\frac{\sqrt{(\kappa - \kappa_i)^2 + (\tau - \tau_i)^2}}{h(\kappa_i, \tau_i)}\right). \tag{1}$$

Here $D$ is a positive-definite function of the distance of a pair $(\kappa, \tau)$ from a particular data point $(\kappa_i, \tau_i)$ and $h$ is a smoothing parameter which determines the "width" of $D$ (which is typically localized in that it decays rapidly when the argument is large). Mathematica's *KernelMixtureDistribution* function was used (7) with a Gaussian Kernel and default bandwidth $h = 0.005$ (see Figure 6(a)). When sampled it produces a distribution which overlaps significantly with the original data (Figure 6(b)). This P.D.F generation procedure was performed for the $\alpha$-helical data. For $\beta$-Strand and Linker data, P.D.F generation was performed separately for the mixed, $\alpha$. left-$\tau$ and right-$\tau$ data sets.

These P.D.F's can be used to generate individual sections of the polypeptide chain, but it is still necessary to link these sections to form a complete chain. The relative orientation of neighboring sections is an extra degree of freedom for the chain's geometry. It is necessary to constrain this freedom empirically. The idea is that, given three points of the previous section

$(m-1)$, some joining $(\kappa, \tau)$ distribution should be used to determine the first point of the next section $(m)$. To do so quadruplets which contained the first $C^{\alpha}$of one section (say a linker) and the last three $C^{\alpha}$'s of the previous section (say an alpha helix), were selected. This was done for each section pair in all molecules and then split into categories linker-helix, helix linker and the same for strands. Then $\kappa$-$\tau$ PDF's were obtained for each type using the same Kernel smoothing technique as in the last section.

##### 1.6 Generating model polypeptide chains.

The algorithm for generating a model backbone, named the CB algorithm (constrained backbone algorithm), is summarized in the flow chart displayed in Figure 7. Its main use here is to generate starting points for model optimisation. Its is assume there are $n$ section labels $l_i$, drawn from the set $\{helix, strand, linker\}$, each identifying the secondary structural nature of a section of length $m_i$.

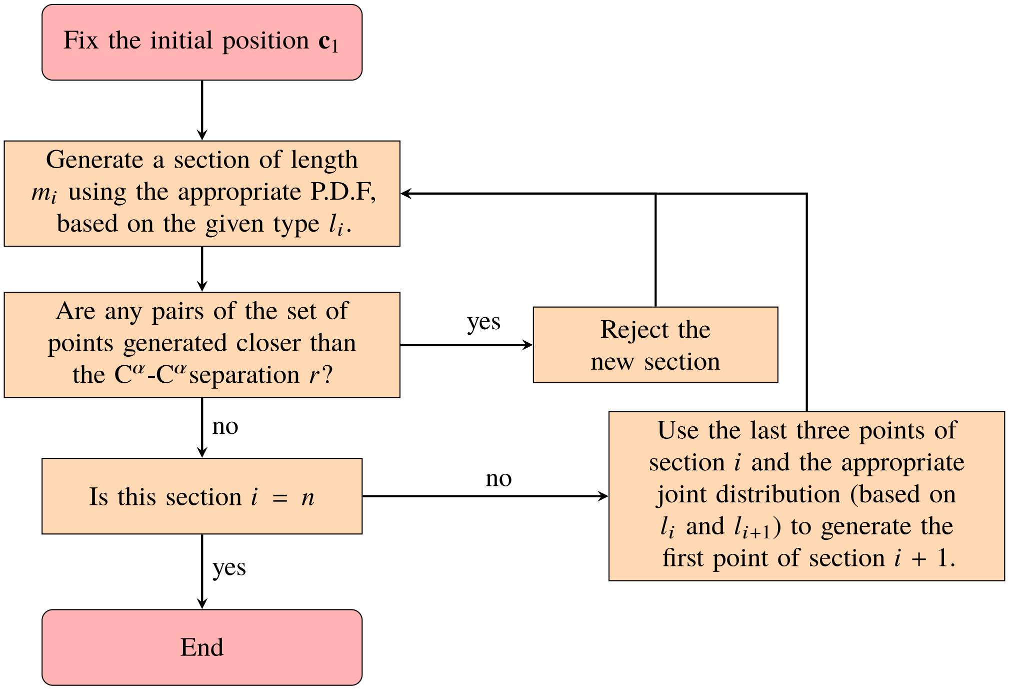


Figure 7: A flow chart describing the CB algorithm for generating model $C^{\alpha}$backbones based on the empirical $(\kappa, \tau)$ distributions obtained for each section type ($\alpha$-helix, $\beta$-strand and linker).

It has been translated into C++. The degrees of geometrical freedom in this structure come from the random generation of the sections and also of relative hinge geometry between sections. The part of the algorithm which rejects sections based on any pairs of points within the section coming too close is to ensures unrealistically folded sectional geometry. The algorithm does not prevent non-local overlap of distinct sections, a constraint which must be additionally enforced, either as an energy penalty in some optimization routine, or by continually re-adjusting individual sections.

#### 2 GENERATING THE HYDRATION LAYER

As all proteins are surrounded by water molecules in solution a hydration layer needs to be added before calculating the small angle X-ray scattering curve. In this section a simple and reliable explicit hydration shell model for an arbitrary backbone curve $\{\mathbf{c}\}_{\mathbf{i}=1}^{\mathbf{n}}$ is detailed. It is empirically calibrated against experimental data.

In this hydration layer description any alpha helical, left $\tau$ or right $\tau$ sections, those with consistent linear geometry, are treated as a whole section; whilst mixed linker and strand sections are decomposed into their individual pairs. Individual sections in this context are referred to as hydration sections, to distinguish from the notion of an individual section in backbone description where mixed linker (and strand) sections are treated as a single unit. It is assumed there are $n_h$ such sections for a chain with $n$ secondary structure units.

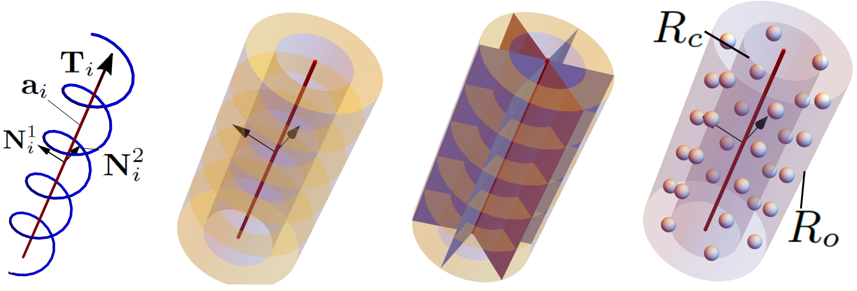


Figure 8: Visualisations of the solvent layer model. (a) the axis curve $\mathbf{a}_i$ at the centre of its an idealised helical section $i$, also shown are the vectors $(\mathbf{T}_i, \mathbf{N}_1^i, \mathbf{N}_2^i)$ which form an orthonormal frame (b). The annular sections (3) formed by splitting the hollow cylinder surrounding $\mathbf{a}_i$. (c) the azimuthal splitting of the sections in (b) which create individual "cells" for the solvent molecules. (d) the points at the centre of the cells of (c), these points are used as the locations of the solvent molecules.

#### 2.1 The initial solvent layer

For each hydration section $i$ a central axis vector $\mathbf{T}_i$ of unit length can be defined. For mixed pairs $(\mathbf{c}_i, \mathbf{c}_{i+1})$ this is just the normalized vector in the direction $\mathbf{c}_{i+1} - \mathbf{c}_i$. For linear hydration sections this is defined by obtaining the central axis of an idealized helix as shown in Figure 8(a). Each section is assigned a central point $\mathbf{c}_i^m$ and length $L_i$ which defines a central axis curve $\mathbf{a}_i(z)$ for the hydration section $i$, as shown in Figure 8(a), and parameterised as

$$\mathbf{a}_i(z) = \mathbf{c}_i^m + \mathbf{T}_i^c z, \quad z \in [-L_i/2, L_i/2]. \tag{2}$$

A unit vector $\mathbf{N}_i^1$, normal to $\mathbf{T}_i$, is used to define an orthonormal frame $(\mathbf{T}_i, \mathbf{N}_i^1, \mathbf{N}_i^2)$ with $\mathbf{N}_i^2 = \mathbf{T}_i \times \mathbf{N}_i^1$. Using this curve $N_{ax}^i$ annular segments can be defined by $N_{ax}^i + 1$ points along the length of the line $\mathbf{c}_i$, with a segment $j$ given by the the equation

$$\begin{aligned} &\mathbf{a}_i((j-1)w + rw - L_i/2) + \rho(\mathbf{N}_i^1 \cos\phi + \mathbf{N}_i^2 \sin\phi), \\ &w = \frac{L_i}{N_{ax}^i}, r \in [0,1],\ \rho \in [R_c, R_o], \phi \in [0, 2\pi),\ j = 0, 1, \ldots N_{ax}^i. \end{aligned} \tag{3}$$

Here $R_c$ is the radius of the inner cylinder where no solvent molecules should be allowed (the "core") and $R_o$ is the outer radius. The annular region between $R_c$ and $R_o$ is where hydration layer molecules can reside. Each annular section is then split into a set of $N_\phi$ "cells" as shown in Figure 8(c). The centre of these cells, which lie on a cylinder of radius $(R_c + R_o)/2$, are chosen to be the central coordinates for the solvent molecules (Figure 8(d)). Thus the set $\mathcal{S}_i$ of solvent centre coordinates is

$$\mathcal{S}_i = \left\{ \mathbf{a}_i\left((j-1/2)\frac{L_i}{N_{ax}} - \frac{L_i}{2}\right) + \frac{(R_c+R_o)}{2}\left[\mathbf{N}_i^1 \cos\left(\frac{2\pi k}{N_\phi}\right) + \mathbf{N}_i^2 \sin\left(\frac{2\pi k}{N_\phi}\right)\right] \middle| \{j\}_1^{N_{ax}},\ \{k\}_0^{N_\phi - 1} \right\}. \tag{4}$$

(Figure 8(d)). This set will also be associated with an $N_\phi$ by $N_{ax}^i$ matrix $S_{kj}^i$ whose entries are either 1, if the solvent is permitted, or 0 if it has been ruled out due to overlapping with some other section of the molecule. So the solvent layer $\mathcal{S}$ is defined by set of pairs $\left\{\mathcal{S}_i, S_{kj}^i\right\}_{i=1}^{n_h}$. $N_\phi$ will not vary section by section but $N_{ax}^i$ will be made to vary with section length.

#### 2.2 Solvent overlap

Once the backbone model has been surrounded by solvent molecules a fast algorithm has been developed to remove overlapping and chemically forbidden solvent molecules form this initial shell. To check for overlap the following steps are taken. Consider the solvent molecule centre $\mathbf{s}_i^{\alpha\beta}$, belonging to hydration section $i$, with $\alpha \in \left\{1, \ldots N_{ax}^i\right\}$ the index along the axis and $\beta \in \left\{0, \ldots N_\phi - 1\right\}$ the azimuthal labelling. To check if $\mathbf{s}_i^{\alpha\beta}$ is either shared or within the forbidden core of section $j$ one must

1. Check if the minimum distance $||\mathbf{s}_i^{\alpha\beta} - \mathbf{a}_j(l)||$, with $l$ the parameter as which this minimum occurs, is either less than $R_c$ (it is forbidden) or between $R_c$ and $R_o$ (shared).

2. Check if the parameter $l$ minimizing $||\mathbf{s}_i^{\alpha\beta} - \mathbf{a}_j(l)||$ is in $[-L_j/2, L_j/2]$.

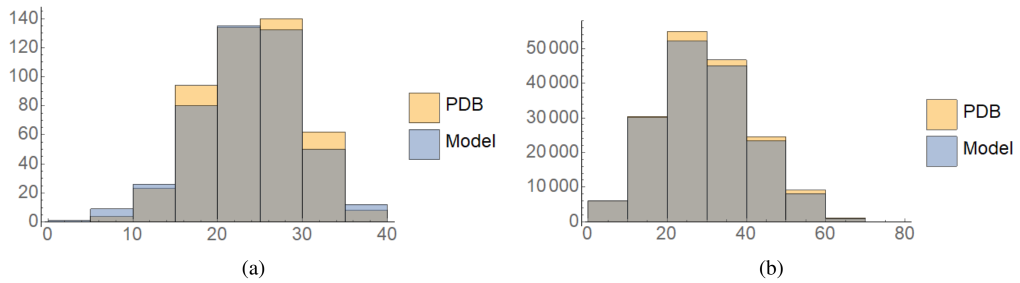


Figure 9: Comparisons of crystallographic and model solvent positions from the crystal structure of a phosphate binding protein PDB=4F1V (8), shown in Figure 4 of the main text.(a) A histogram comparison of the distances of the solvent molecules from the centre of mass of the backbone for the two distributions shown in (a) and (b) of the man text figure (in angstroms). The model distribution is slightly more evenly spread. (b) A histogram of the solvent-solvent distribution distances (in angstroms) of the two sets of solvent data, the two histograms are very similar.

Minimising $||\mathbf{s}_i - \mathbf{a}_j(l)||$ gives

$$l = (\mathbf{s}_i^{\alpha\beta} - \mathbf{c}_j^m) \cdot \mathbf{T}_j. \tag{5}$$

In what follows the minimum distance, given by inserting (5) into $||\mathbf{s}_i^{\alpha\beta} - \mathbf{a}_j(l)||$, is labelled $d_{ij}^{\alpha\beta}$.

#### 2.3 Checking overlap for the full molecule

In order to minimize the number of calculations required to generate the hydration shell the following protocol has been developed.

1. Initialise a set of matrices

$$\left\{S^i_{\alpha\beta} | \, \alpha_i \in \left\{1, \dots N^i_{ax}\right\}, \beta_i \in \left\{0, \dots N_\phi - 1\right\}, \right. \\ \left. i \in \{1, \dots n_h\}\right\}, \tag{6}$$

   all of whose entries are 1. Here $n_h$ is the number of hydration sections of the backbone curve. As discussed above the entires of these matrices indicate if the solvent is in the final hydration shell or not. Thus initially we assume they are all part of the shell.

2. Check any of the solvent molecules of section 1 lie in the hydration layer of section 2 ($R_c \leq d_{12}^{\alpha\beta} \leq R_o$). If this inequality is satisfied then switch the 1 at index $(\alpha, \beta)$ of matrix $S^1_{\alpha\beta}$ to a zero as the solvent molecule is shared.

3. Check if any solvent molecules of section 2 lie in the forbidden tubular neighborhood of section 1, ($0 \leq d_{21}^{\alpha\beta} \leq R_c$). If this inequality is satisfied then switch the 1 at index $(\alpha, \beta)$ of matrix $S^2_{\alpha\beta}$ to a zero as the solvent molecule is forbidden.

4. Repeat steps 2, and 3 for section 1 with all $j \in \{3, \dots n_h\}$ for all molecules whose index $(\alpha, \beta)$ in the updated $S^i_{\alpha\beta}$ is non zero, *i.e.* check the inequality ($R_c \leq d_{1j}^{\alpha\beta} \leq R_o$)

5. Repeat 4 for section $j \in \{3, \dots n_h\}$ with section 1, i.e. check $0 \leq d_{j1}^{\alpha\beta} \leq R_c$.

6. Repeat steps 2-5 with section 2 and all $j \in \{3, \dots n_h\}$, being careful to use the updated matrix $S^i_{\alpha\beta}$, which may have lost some solvent molecules overlapping with the core of 1.

7. Repeat step 6 sequentially on all $i \in \{3, \dots n_h - 1\}$ for al $j \in \{i+1, \dots n_h\}$, always taking care to update the state of all matrices.

Once this last step has been completed the matrix $S^i_{kj}$ is used as a lookup table for the existence of a given solvent molecule for the purpose of calculating the scattering. This algorithm was translated into a C++ code.

#### 2.4 Hydration layer parameters

In addition to the shape of the backbone $\{\mathbf{c}_i\}_{i=1}^n$ the hydration layer model requires values for four other parameters $(R_c, R_o, N_\phi, r)$, where $r$ defines a ratio of the number of C$^\alpha$'s in a section to the number of solvent rings through $N_{ax}^i = \text{round}(m_i r)$ (round up to the nearest whole number based on the number of C$^\alpha$'s in the section). Suitable fixed values for these constants were determined as follows. The positions of solvent molecules were extracted from ultra high resolution crystallographic data in three PDB files 4F1V (Periplasmic phosphate binding protein (8)), 1US0 (human Aldose Reductase (9)) and 2VB1 (Lysozyme (10)). The number of solvent molecules were adjusted to account for the solvent occupancy and $\beta$ values to give a ratio of solvent molecules to C$^\alpha$'s in the molecule of 1.25, this is approximately in-line with water molecules detected in the first hydration shell (11). Using this data two histogram plots for each molecule were compiled, one of the set of distances of the solvent molecules from the molecule's centre of mass and one of all solvent-solvent distances. The first gives some indication of the distribution both within and surrounding the molecule, the second gives some idea of the spread of solvent distances. Manually varying the set $(R_c, R_o, N_\phi, r)$ the parameter values (6Å, 7Å, 6, 1) were found to give the best fit to all three data sets. For example Figure 4 of the main text depicts the crystallographic (a) and model distributions (b) for the 4F1V structure, the model distribution is slightly more evenly distributed. Comparisons of the two histograms for the 4F1V solvent positions and the model hydration layer are shown in Figure 9(a) and (b) and are similar.

### 3 DETERMINATION OF THE PARAMETERS OF THE SCATTERING MODEL

#### 3.1 Amino acid form factors

The form factors $f_{am}$ for an amino acid whose scattering is centered on its C$^\alpha$position is

$$f_{am}(q) = f_b(q) - \rho_{ex} f_{ex}(q), \tag{7}$$

where $f_b$ is the scattering of the amino acid in a vacuum and $f_{ex}$ is the adjustment due to the excluded volume of solvent in the comparison scattering data and $\rho_{ex}$ a constant mondulating its effect. Whilst $f_b(q)$ of individual amino acids are known, an average representation is used to significantly reduce the time taken to evaluate the Debye formula. The distances $r_{ij}$ are separated into bins of fixed width, then, assuming the amino acids all have the same form factor $f_b$, the contribution to the Debye formula from all the distances in a given bin can be approximated as a single calculation. A bin size of 0.2Å was found to give sufficient accuracy for $q \in [0, 0.4]$ (the range used in the fitting algorithm). To represent this averaged amino acid form factor $f_b$ an empirically determined model is utilized with a five-factor exponential representation

$$f_b = \sum_{i=1}^{5} A_i \mathrm{e}^{-B_i q^2} + C, \tag{8}$$

where $\{A_i, B_i\}_{i=1}^5$ and $C$ constants. This is the same mathematical form as used to fit to individual amino acid form factors (12).

The excluded volume effect is captured using an exponential model in the form

$$f_{ex}^a(r_w, q) = v(r_w)\mathrm{e}^{-\pi q^2 v(r_w)^{3/2}}, \quad v(r_w) = \frac{4\pi}{3} r_w^3, \tag{9}$$

where $r_w$ is the average atomic radius of the atom (13–15). To calculate the excluded volume for amino acids PDB coordinates for all 20 amino acids (16), and values of $r_w$ for Carbon, Nitrogen, Oxygen, Hydrogen and Sulphur (*e.g.* (17)) were used to compute the excluded volume scattering centered at the C$^\alpha$through

$$f_{ex}^{am}(q) = \sum_{i=1}^{N_{am}} f_{ex}^a(r_{wi}, q) \frac{\sin(q r_i^\alpha)}{q r_i^\alpha}, \tag{10}$$

where $r_i^\alpha$ is the distance of molecule $i$ from the C$^\alpha$molecule and $N_{am}$ the number of molecules in the amino acid. Since $f_b$ does not discriminate individual amino acids this value $f_{ex}^{am}$ was averaged over all 20 amino acids, weighted by their abundance in globular proteins (see (18)). This averaged function, shown in Figure 10(a), gives $f_{ex}(q)$. Finally (7) includes a constant $\rho_{ex}$ which modulates the effect of the excluded volume scatter by comparison to $f_b$, this value is constrained to lie withing 0.75 and 1.25 (similar constraints are used in (13–15)).

The abundance weighted averaging procedure could also have been used to calculate an average scattering function for the amino acids, this function is labelled $\hat{f}_b(q)$ here and it is shown in Figure 10(a). In practice the extra freedom offered by the empirical law (8) lead to better results, however, as will be discussed shortly, this average $\hat{f}_b(q)$ was used to constrain the values of the constants and $\{A_i, B_i\}_{i=1}^5$ and $C$ so that the function $f_b(q)$ remained physically plausible.

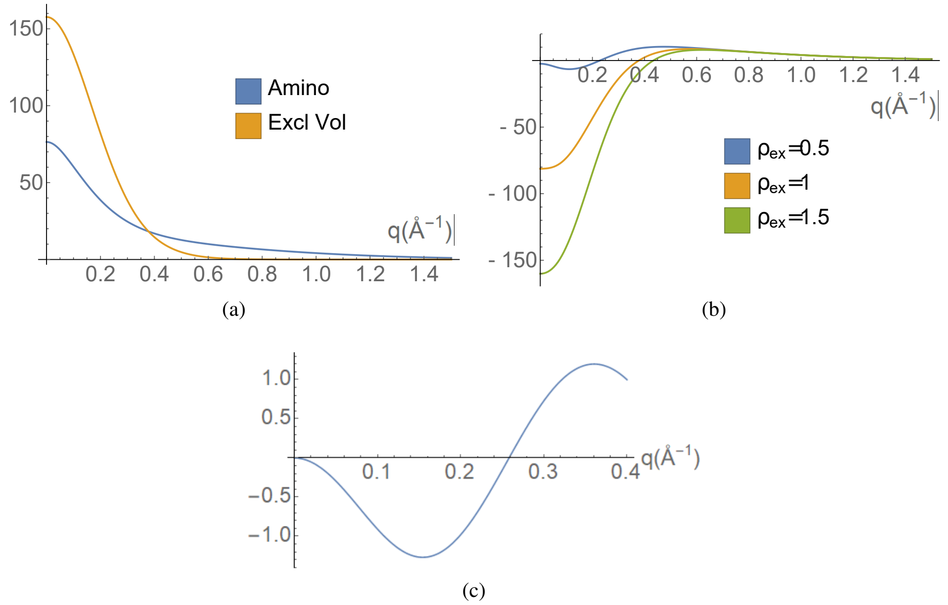


Figure 10: Representative curves associated with the scattering model. (a) the excluded volume scattering $f_{ex}(q)$, calculated using (10) and averaged by abundance per amino acid. Also shown in $\hat{f}_b(q)$ the abundance weighted amino acid scattering. (b) the difference between the averaged amino acid scattering $\hat{f}_b$ and the excluded volume scattering for various values of $\rho_{ex}$, the physically acceptable range is constrained to lie between 0.75 and 1.25, similar to (13–15). (c) A scattering function $f_{am}(q)$ obtained by minimizing (12) which was rejected as non-physical, its morphology is similar to the $\rho_{ex} = 0.5$ case which is outisde the physically expected range.

##### 3.2 Hydration layer form factors

The scattering form for an individual water molecule in the hydration layer is

$$f_h(q) = \rho_h(2 f_{hy}(q) + f_{ox}(q)), \tag{11}$$

where $f_{hy}$ and $f_{ox}$ are the vacuum scattering of Hydrogen and Oxygen respectively (there are highly accurate Gaussian approximations for these functions (12)). The constant $\rho_h$ is required to adjust the scattering amplitude to represent the hydration effect, it is to be empirically determined (as in (15)).

##### 3.3 Determining the scattering parameters

To determine the constants $\{A_i, B_i\}_{i=1}^{5}, C, \rho_{ex}, \rho_h$ a set of well resolved and reliable PDB structures for which high quality scattering data was available were selected, namely 1LYZ (Lysozyme (19)), 1HRC (Horse heart cytochrome (20)), 1C0B (Bovine erythrocyte enzyme superoxide dismutase (21)), 3V03 (Bovine serum albumin (22)), all selected from the SASBDB repository (23). The PDB C$^{\alpha}$coordinates were taken as the molecule backbone, then the hydration layer model described in the previous section was constructed yielding coordinates for the hydration layer solvent molecules. Using these coordinates and the Debye scatting formula with (7) and (11), parameter dependent model scattering curves $I_m\left[\{A_i, B_i\}_{i=1}^{5}, C, \rho_{ex}, \rho_h\right](q)$ were compared to this experimental data.

##### 3.4 Smoothing the experimental data $I_e(q)$

A number of different approaches have been proposed to filtering experimental noise (*e.g.* (13, 15, 24)). Here we adopt the recent approach of Rambo and Tainer was used as it was shown to reduce model fitting error((24)). The data is smoothed by separating it into $n_s$ equal size bins, with $n_s$ determined by the molecule's size using the Shannon sampling theorem. If $d_{max}$ is the maximum value of the pair-pair distances of all molecules in the protein then $n_s = \text{round}(d_{max} q_{max}/\pi)$, where $q_{max}$ is the maximum value of the momentum transfer in the scattering data used. Values from this each bin are selected in some fashion (the median of a set of random samples for example) . The smoothed curve obtained from this procedure is labelled $I_e^s(q)$.

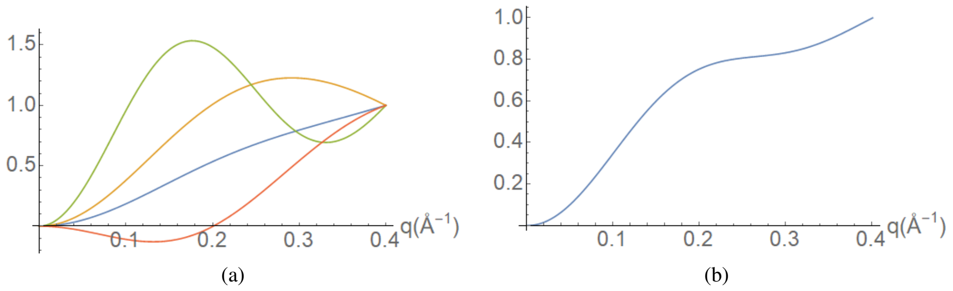


Figure 11: The scaled amino acid scattering curves $f_{am}(q)$, obtained from the scattering data fits shown in Figure 5 of the main text. (a) the various $f_{am}(q)$ curves terms for each case shown in Figure 5 (of the main text). (b) the average of the four curves shown in (a).

##### 3.5 Model optimization against the experimental BioSAXS data

Quality of the model fit was determined by the following $\chi$-squared function

$$\chi_s^2 = \frac{1}{n_s - 1}\sum_{i=1}^{n_s}\left[\log(I_m(q)) - a - \log(I_e^s(q))\right]^2, \tag{12}$$

where $a$ is an adjustment parameter used to account for the unknown concentration of the protein. Mathematica's *NMinimize[ ]* routine was used to minimize over the set $\{A_i, B_i\}_{i=1}^5, C, \rho_{ex}, \rho_h, a$ (7). A number of additional constraints were applied. First that the zero scattering values of the amino acid scattering $f_{am}(0)$ (given by (8) and (10)) and the hydration scattering $f_h$, given by (11) must always differ by the same value. This constraint was imposed in order to keep the relative effect of the hydration layer and amino acid scattering consistent between molecules. A sensible value for this fixed difference is the difference between the abundance average amino acid scattering $\hat{f}_b$ and $f_h(0)$. Thus the hydration scattering is given by a function $f_{ha}$:

$$f_{ha}(q) = f_h(q) + f_{am}(0) - \hat{f}_{am}(0). \tag{13}$$

The second constraint was imposed owing to the fact that some fits produced a function $f_{am}(q)$ which had a physically unrealistic form. The function $\hat{f}_{am}(q) = \hat{f}_b - \rho_{ex} f_{ex}(q)$ shown in Figure 10(b) for various values of $\rho_{ex}$ was used as a guide. The basic morphology is clear, one should expect a shape which might look somewhat like a sigmoid. Forms of $f_{am}(q)$, obtained by minimizing $\chi_s^2$ were rejected if they differed too much form this sigmoidal shape, an example of a rejected form is shown in Figure 10(c).

The various fits of examples taken from the SAS database are shown in Figure 5 of the main text all with a significantly low value $\chi_s$ value. The scattering functions $f_{am}(q)$ are shown (scaled) in Figure 11(a).

###### 3.5.1 Averaged form functions

The actual protein backbones used are amongst the set of structures which could be generated by the CB model generation algorithm described in (Section 1.5), so it is plausible that the model could be used in an ab-inito setting to generate this structure. However the final ab-initio optimization will not be simultaneously fitting both model parameters ($C^\alpha$positions $\mathbf{c}_i$) and scattering parameters, this would run the risk of over-fitting and potentially promote unreasonable structures, so the parameters obtained for each fit must in some way be combined to obtain a fixed set $\{A_i, B_i\}_{i=1}^5, C, \rho_{ex}, \rho_h$.

To this aim the average of the scattering formulae used to obtain the fits shown in Figure 5 (of the main text) was calculated by mutually scaling the scattering formulae between 0 and 1. The scaled curves are shown in Figure 11(a), the average, shown in Figure 11(b), has a monotonic rise but more variation than the average $\hat{f}_{am}(q)$ (shown in Figure 10(a)).

#### 4 EVALUATING STRUCTURAL SIMILARITY

##### 4.1 The need for a knot based structure comparison

In order to compare the similarity of folding of two curves we developed a novel technique that enables us to:

1. Validate of the fitting algorithm's predictions on reliable structures taken from the PDB *i.e.* those such as Lysozyme with high quality SAXS data.

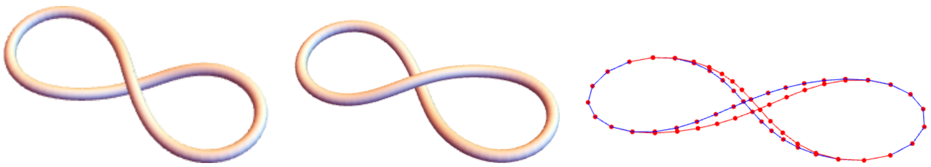

(a) (b) (c)

Figure 12: *Figure 8* curves. (a) and (b) are *figure 8* curves which can be inter-converted by applying a half twist to the curve about its centre. (c) are imagined discretizations of the two curves, from which RMSD measure can be calculated.

2. Classify predictions. A reasonable procedure would be to run the algorithm numerous times and compare the multiple predictions. It would be beneficial to be able to discern which predictions are "essentially the same" in that they only differ by small local deformations (as one might expect in solution).

The conventional $C^{\alpha}$backbone structural comparison techniques are based on root mean squared deviations (RMSD) of paired distances of the $C^{\alpha}$coordinates (25). The basic idea is one structure is superimposed on the other in order to minimize the root mean square distance between points at the same index along the chain. As discussed in (25) this approach suffers from some significant drawbacks. Firstly, the superimposition process is an ambiguous, with multiple possible solutions. Secondly a strong difference in a single loop can lead to large deviations. Thirdly, the flexibility of linker sections can lead to significant ambiguity, both in the model prediction and any comparative crystallographic structures. There are various ways of trying to minimize these problems, such as weighting the distance error estimate to limit the effects of more flexible units or finding the maximum chain length with a small RMSD, but many of the above problems still persist. Other more subtle variations on this superimposition approach exist (25). However, for the purpose of this ab-initio fitting there is ambiguity, both in the experimental scattering data, due to the random molecule orientation, and the temporal flexibility of a structure in solution. So even if the predicted structure folded in approximately the right manner, one should expect a comparison between two structures or with a PDB structure could lead to a significant build up of small errors, unlike in crystal structure comparison where the difference between structures often only involves a difference in a small subsection of the structure (*e.g.* (26)). One other notable alternative to shape comparison was proposed by Røgen and Bohr, based on higher order Gauss integrals (27). As noted by the authors the majority of the quantities do not have an obvious physical interpretation and this approach was not pursued here.

##### 4.1.1 Topological distance

In addition to the issues mentioned above, the view was taken that the distance based approach would be prone to more significant problems, as now discussed. As an extreme idealization of a closed polypeptide chain consider two *figure 8* curves created by twisting a circle in a right handed or left handed fashion until the curve is close to figure 8 contact, as shown Figures 12 (a) and (b). There are two ways to turn the two curves into each other, one is to apply a half twist about the centre of the curve to create the *figure 8* curve of opposite chirality. For such a change in conformation the distance travelled by all points of the curve (as discretized in Figure 12(c)) would be significant. On the other hand by passing the curve through itself at the point of closest approach a right-handed figure 8 can be turned into a left-handed one by only moving a few points by a very small distance. This is essentially the distance which would be measured by a root mean squared error measure such as

$$D_{rmsd} = \frac{2}{(L_1 + L_2)}\sqrt{\sum_{i=1}^{n} d_i}, \tag{14}$$

where the $d_i$ are the distance between points at index $i$ on both curves, and $2/(L_1 + L_2)$ the mean length of the two curves. The value of $D_{rmsd}$ is about 20 percent, such figure would be considered structurally similar by distance based criteria (see e.g.(25)). This measure, however, is less than ideal because it is not representative of the average "distance" each point would have to move to turn one structure into the other in solution without cutting (*i.e.* by a full rotation). It is clear some other measure of the structure's similarity is necessary.

As a second example consider the trefoil curve shown in Figure 13(a). Two further curves, Figures 13(b) and (c) are created by adding random disturbances to the structure. Each point $\mathbf{c}_i$ is altered by adding random perturbations *i.e.* $\mathbf{c}_i \to \mathbf{c}_i + (U(0,r), U(0,r), U(0,r))$, with $U(0,r)$ a uniform distribution of numbers between 0 and $r$. For $r = 1$ (Figure 13(b)) the curve is still clearly a trefoil, *i.e.* its fold is essentially the same as the original curve. For $r = 3$ ( Figure 13(c)) the curve has become significantly more complex in its entanglement. The $D_{rmsd}$ measures for each of (b) and (c) in comparison to (a) are 16.8% and 17.67% respectively. So despite being essentially different folds the RMSD measure registers very little difference.

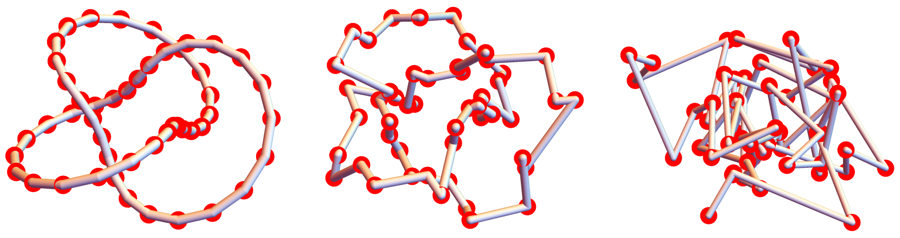


Figure 13: A trefoil curve and distorted variants of the structure. (a) a discretized trefoil curve. (b) a structure obtained from (a) by applying random displacements to each point of (a) within a radius of $\sqrt{3}$. The "knotted" entanglement of (a) is still clearly visible. (c) a structure obtained from (a) by applying random displacements to each point of (a) within a radius of $3\sqrt{3}$. Its entanglement is dramatically different from (a) and (b).

Of course these examples are a significant idealization but intuitively one should expect this problem to become more significant for more complex structures. To define an alternative measure of curve similarity techniques from knot theory were adopted.

#### 4.2 Secondary knot similarity

Knot topology is the field of mathematics which can give an answer to the question "can one curve be turned into another without self-intersection? (see *e.g.* (28) for an introduction to the subject). It typically concerns closed curves, curves with no end points. The *unknot* is any curve which can be deformed into a circle without crossing itself. By contrast knots are curves which **must** be cut in order to be turned into a circle. Knots can then be classified in various ways into curves which can be deformed into each other without self-crossing.

This can only be a precise question if the curves are closed as one can always untangle an open ended curve. To attempt classification of open curves they are transformed into closed curves by joining the ends of the open curve. But then the choice of closure can affect the type of curve. In order circumvent this problem, the following procedure to define the *knot fingerprint* of any given polypeptide chain was adopted from (29):

1. Surround the backbone $\{\mathbf{c}_i\}_{i=1}^{n}$, represented by its $C^{\alpha}$positions, with a sphere.
2. Take a subsection of the backbone $\{\mathbf{c}_i\}_{i=j}^{k}$ $(j < k)$. Randomly select two points on the sphere and join the subsection ends $\mathbf{c}_j$ and $\mathbf{c}_k$ to these points. Create a close curve with a geodesic lines on the sphere.
3. Classify the knot (the closed curve) using some technique from the knot theory literature (see e.g. (28)). This involves simplifying the curve whilst maintaining its topology.
4. Repeat this process with random closures some number $N$ times (10000 in this study) collecting a list of the knots produced.
5. Select the most commonly occurring knot type from the $N$ closed curves.
6. Repeat this for all $j, k \in [1, n]$ such that $j < k$ and $k - j > 3$ (four points are necessary to create a discrete knot).

One can then plot this data on a staircase chart with $j$ and $k$ on the axes and each square of the domain coloured by its most common knot (similar to Figure 14(c)). This diagram is the knot fingerprint. The Knot Prot database utlizes this procedure to calculate the fingerprint which is found to be preserved across protein families (30), even when there is low sequence identity (29). For the purpose of this study a C++ code was developed to perform this fingerprint generation, as the aim and analysis of its output differ from those of the Knot Prot data base.

In this study the knot identification part of the task is performed by a code written by Dr Mark Miller which evaluates the Jones polynomial of the given closed curve, deduced from a projection this curve. The main computational expense is associated with evaluating the number of unkots that result from all possible combinations of splits of the crossings. To improve the efficiency of the calculation, the number of crossings is reduced before the splitting analysis in two steps. Firstly, projections in several random directions are compared and the projection with fewest crossings is selected. Secondly, Reidemeister type-1

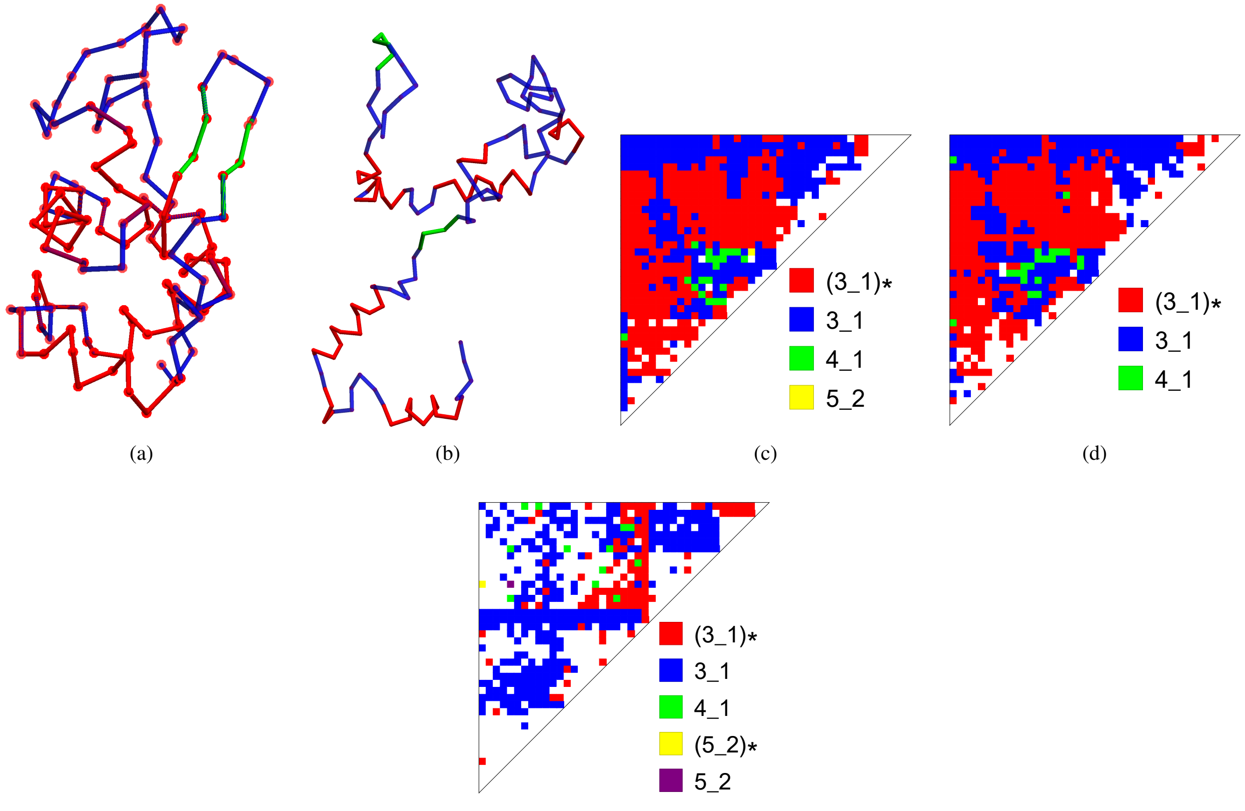


Figure 14: $C^{\alpha}$traces of Lysozyme models and their secondary knot fingerprints. (a) the backbone curve of the 1LYZ pdb file. The $\alpha$-helices are shown in red, strand structures green, and linker sections blue. (b) a randomly generated structure based on secondary structure sequence of the 1LYZ pdb, generated using the CB algorithm. (c) the secondary knot Fingerprint for the 1LYZ backbone shown in (a). The knot types are labelled according to their Rolfsen classification (33). The trefoil knots of both handedness $3_1$ and $(3_1)_*$ dominate. (d) the secondary knot fingerprint for the 1AKI backbone structure (also Lysozyme) (d). (e) the secondary fingerprint for the structure shown in (b), it differs significantly from (c) and (d) and has a larger range of knots present.

moves are used to remove any trivial loops. The routines are adapted from code originally used to identify knots in chains of dipolar colloidal particles (31, 32).

The interest here is to discriminate structures which have the same knot finger print, but differing fold geometry. Lysozyme for example (whose backbone is shown in Figure 14) has a staircase diagram for which each square is the unknot. Just generating a random structure with the same set of secondary elements as Lysozyme (using the technique described in Section 1.5) tends to lead to curves with the same fingerprint. But these structures differ significantly, for example the random structures are often far less compact (*c.f.* Figure 14(a) and (b)) even though their staircase diagrams are identical.

If, however, the fingerprint structure was instead constructed with each square being labelled by the **second** most common knot then the "*secondary finger prints*" differ significantly for the two curves shown in Figure 14(a) and (b) (*c.f.* Figure 14(c) and Figure 14(e)). In this case the majority of the squares indicate the second most common knots are Trefoil knots (left and right handed), the simplest knot which only requires one cut to be undone. By comparison the secondary knot fingerprint for a different set of coordinates of the same protein (Lysozyme, PDB=1AKI (34)), shown in Figure 14(d), is significantly similar to the original 1LYZ crystal structure's (secondary) finger print. This indicates that secondary knot information could be used as a comparison of fold similarity (if the two primary knot finger prints match). Indeed one might go to lower levels *i.e.* tertiary and quaternary fingerprints (although in this study the first two are found to suffice). To this aim a quantity measuring the similarity $l^{th}$ most common knot fingerprint, called the "*knot fingerprint statistic*" was defined.

###### 4.2.1 The knot fingerprint statistic

The knot fingerprint statistic $\mathcal{K}_l$ yields a number between 0 and 1, with 1 implying the two knot fingerprints of interest are identical on a square-by-square basis, and 0 if they are completely dissimilar. It accounts for the number of knots in each square of the $l^{th}$ fingerprint and the similarity in number (per square) between the two ($l^{th}$ level) fingerprints, a value of 1 implies these values are identical for all squares.

Consider two discrete curves $K_1 = \left\{\mathbf{c}^1\right\}_i^n$ and $K_2 = \left\{\mathbf{c}^2\right\}_i^n$. For each subsection $\left\{\mathbf{c}_i^1 | \, i \in [j,k]\right\}$, $\left\{\mathbf{c}_i^2 | \, i \in [j,k]\right\}$ label $i_1^{jk}, i_2^{jk}$ as the knot type of the $l^{th}$ most common knot and $n_1^{jk}, n_2^{jk}$ the number of times that it occurs. Define a "weight function" $w_{jk} \in [0,1]$,

$$w_{jk} = \exp\left[\ln 2\left(\frac{n_1^{jk} + n_2^{jk}}{S_{max}}\right)^p\right] - 1, \tag{15}$$

$$S_{max} = \max_{j,k \in [1,n],\, k-j>3} n_1^{jk} + n_2^{jk}. \tag{16}$$

An exponentially increasing weight is assigned based on the relative number of knots produced for a given subsection of $K_1$ and $K_2$ (relative to the maximum number over all subsections $S_{max}$). The higher the value of $p$ the greater the relative weight given to larger sums $n_1^{kj} + n_2^{kj}$, all calculations in this note used $p = 5$ which, as will be seen shortly, was found to give a statistic which had excellent properties in distinguishing folds. Using this weighting the following matching function function $c_{jk}$ is defined

$$c_{jk} = \begin{cases} w_{jk}\left(1 - \left|\frac{n_1^{jk} - n_2^{jk}}{n_1^{jk} + n_2^{jk}}\right|\right) & i_1^{jk} = i_2^{jk}, \\ 0 & i_1^{jk} \neq i_2^{jk}. \end{cases} \tag{17}$$

This assigns a weighting if there is a match in knot type and gives preference to matches which have similar numbers $n_1^{kj}, n_2^{kj}$. The knot fingerprint statistic $\mathcal{K}_l(K_1, K_2)$ is following real number which takes values on $[0,1]$

$$\mathcal{K}_l(K_1, K_2) = \frac{\sum_{k=4}^n \sum_{j=1}^{k-3} c_{jk}}{\sum_{k=4}^n \sum_{j=1}^{k-3} w_{jk}}. \tag{18}$$

This can only be one if all $i_1^{jk}, i_2^{jk}$ are the same and have exactly the same individual number of occurrences $n_1^{jk} = n_2^{jk}$.

##### 4.3 Fingerprint statistics as a tool to compare and discriminate protein models.

In the following paragraph the potential of secondary knot fingerprint statistics as a means to compare and discriminate different protein models, we begin by using the hypothetical *figure 8* example.

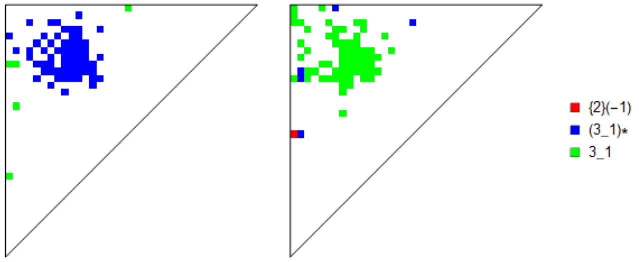


Figure 15: Secondary knot fingerprints for the *figure 8* curves shown in 12(c).

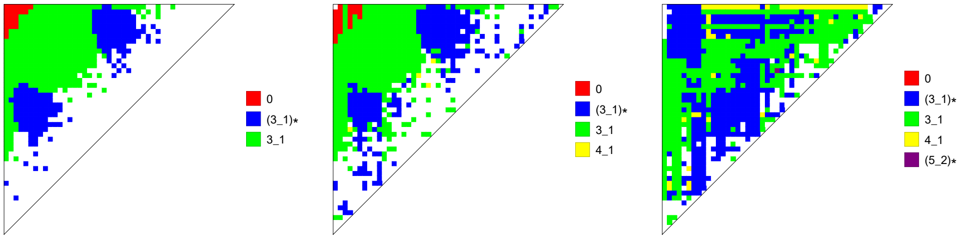


(a) (b) (c)

Figure 16: The secondary knot fingerprints of the curves shown in Figure 13(a)-(c). The labelling (a)-(c) here is the same.

###### 4.3.1 *Figure 8* curves

As one would expect the primary finger prints had the most common knot is the unknot on all squares so it is left to the secondary fingerprint to discriminate. The curves are shown in Figure 12(c) and the secondary knot fingerprints in Figure 15. The secondary fingerprints are dominated by trefoil knots, $3_1$ in the Rolfsen knot classification, these occur in opposing orientations, $3_1$ is right handed and $(3_1)_*$ left-handed, (shown respectively in green and blue in Figure 15). The value of $\mathcal{K}_2$ is zero in this idealized case so the structures are classed as being topologically distinct.

###### 4.3.2 Trefoil curves

The three (secondary) fingerprints for the trefoil derived curves depicted in Figure 13 are shown in Figure 16. As expected the first two curves in Figure 13 (a) and (b) have significantly similar fingerprints, whilst the the finger print of the curve in Figure 13(c) differs significantly. This conclusion is echoed in the secondary finger print statistics. Labelling the curves $K_1$ (a), $K_2$ (b) and $K_3$ (c), it was determined that $\mathcal{K}_2(K_1, K_2) = 0.71$, $\mathcal{K}_2(K_1, K_3) = 0.03$ and $\mathcal{K}_2(K_2, K_3) = 0.03$ (both 0.03). So once again the fingerprint statistic correctly discriminates these structures entanglement where the RMSD measure fails to do so.

###### 4.3.3 Application of the secondary knot fingerprint statistic to protein structures

The coordinates of Lysozyme structures in different space group, *i.e.* in a different local environment, were used to assess the knot fingerprint technique for very similar, yet not identical sets of coordinates. The following PDB entries for Lysozyme were used 1AKI (34), 1HSW (35), 1UC0 (36), 4R0F (37) and 3LZT (38). For comparison 50 random structures were generated using the CB algorithm with the same secondary structure assignment as the 1LYZ pdb file. Example finger prints are shown in Figure 14 (c) and (d) for 1LYZ and 1AKI and (e) for a random structure. As expected most knots are trefoil knots of both chirality, in this case $\mathcal{K} = 0.943$ for the two structures, whilst $\mathcal{K} = 0.132$ for a comparison of Figure 14 (c) and (e). A histogram of the $\mathcal{K}_2$ values of all the random structures and Lysozyme structures (both in comparison to the 1LYZ structure) is shown in Figure 17(a), there is a significant gap between the two sets of data, the mean value of $\mathcal{K}$ of the PDB data set 0.893 and 0.042 the random data set.

###### 4.3.4 The secondary knot fingerprint statistic as a sensitive measure of structural changes.

In order to assess how structural changes of the polypeptide chain manifest in the knot fingerprints, deformed structures were generated by altering $n$ randomly chosen sections of a crystal structure's backbone curve, replacing them with new sections

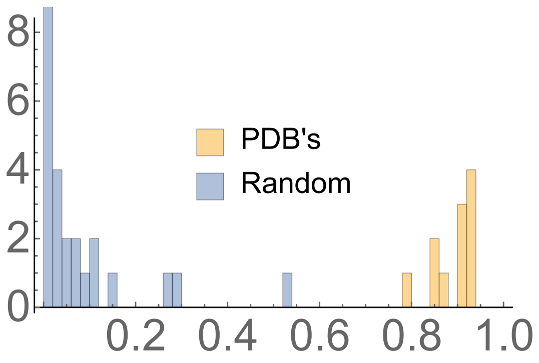


Figure 17: A comparison of the secondary knot statistic $\mathcal{K}_2$ for various Lysozyme PDB structures and randomly generated structures. All statistics are calculated with respect to the 1LYZ PDB coordinates. There is a significant gap between the two data sets indicating $K$ classify the two sets as (generally) significantly different.

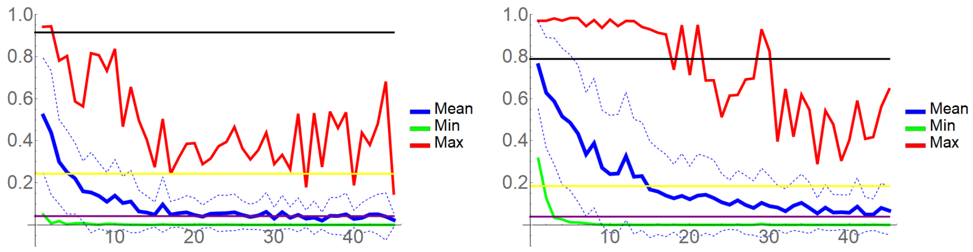


(a) (b)

Figure 18: Plots of $\mathcal{K}_l$ for deformed PDB structures as a function of the number of (randomly) changes sections. The curves shown are the mean value, with dashed limits of 1 S.D, the minimum and maximum values. (a) is for Lysozyme (based on the 1LYZ structure) The horizontal lines are respectively the average of the PDB $\mathcal{K}$ values (black) line 0.915, the ab-initio fit average (yellow), 0.243, and the random structure average (purple) 0.0414. (b) is for Ribonuclease, based on the 1C0B structure.

generated using the appropriate P.D.F to generate sets of $(\kappa, \tau)$ pairs, and using the algorithm detailed in Appendix 1 to construct the new section and link it to the existing structure. Fifty such deformed structures were generated for each number of changes $n$. The effect of these changes on $\mathcal{K}_2(K_{pdb}, K_n)$, with $K_{pdb}$ the original structure and $K_n$ a structure obtained from $K_{pdb}$ by $n$ changes, are summarized in Figure 18(a) for Lysozyme (the PDB 1LYZ) and Figure 18(b) for Ribonuclease (the PDB 1C0B). For Lysozyme, the mean drops rapidly to the same value as the random average after about 15 changes. For comparison the mean $\mathcal{K}_2$ values of alternative Lysozyme PDB's, the SAXS curve optimized Lysozyme structures (see the main paper for details) and the random CB algorithm generated Lysozome structures are shown. Notably the minimum value is always significantly low, even after one change, whilst the maximum value shows a similar (but more variable) drop off to the maximum value found for a randomly generated structure (see Figure 17). For 1C0B the story is similar except the decay in both the mean and maximum values with $n$ is slower. There is some indication the entanglement of the 1C0B structure is more resilient to changes in its individual units than 1LYZ, but in both cases the finger print statistic shows (on average) the expected decay as the structure is increasingly altered.

###### 4.3.5 Comparison between similar sized monomeric proteins

In order to show the general applicability of this structural/curve comparison technique a set of small monomeric proteins were selected for comparative analysis using $\mathcal{K}_2$. Sets of proteins structures of similar molecule weight (number of amino acids) were collected. The sets are

1. 100-125 amino acids, 1C0B (Bovince Pancreantic nuclease (21)), 1HRC (Horse Heart Cytochrome (20)) and 1CMB (E-coli Met Repressor (39)).

2. 150-170 amino acids, 1HL5 (Superoxide Dismutase (40)), 1WLA (Myoglobin (41)), 4UD1 (MERS CoV nucleocapsid (42)).

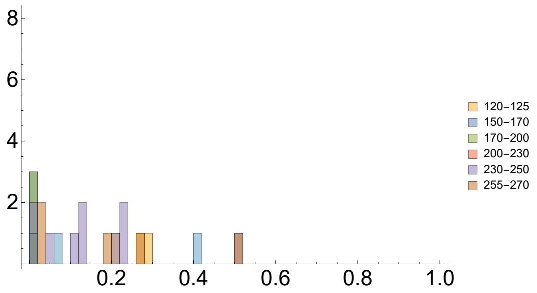


Figure 19: A histogram of of the secondary finger print statistics $\mathcal{K}_2$ for a variety of crystal monomer structures, categorized and compared based on the number of amino acids in the monomer. The various categories and their values are color coded. None show a significant match.

3. 170-200 amino acids, 1IER (Horse Spleen Apoferatin (43)), 3K3K (Abscisic acid (44))
4. 210-230 amino acids , 3PXJ (Abscisic acid receptor pyrabactin resistence PYR1 protein (45)), 4V07 (Protease (46)), 1L6W (Fructose-6-phosphate aldolase (47)), 1NSJ (Isomerase (48)).
5. 230-250 amino acids, 1A53 (Synthase (49)), 1QO2 (ribonucleotid isomerase (50)), 2W6R (Artificial 8-barrel protein (51)), 4PCF (Triosephosphate isomerase (52)).
6. 255-270 amino acids, 1M40 (TEM1 (53)), 1V9E (Bovine Carbonic Anhydrase (54)), 1OK6 (Archaeal fructose (55)).

These structure were compared to each other (pairwise, within categories) using $\mathcal{K}_1$ and $\mathcal{K}_2$.

All primary $\mathcal{K}_1$ measures were above 0.97 so it was left to the secondary knot fingerprint statistic $\mathcal{K}_2$ to distinguish the structures. The results are shown in Figure 19, none are much above $\mathcal{K}_2 = 0.5$, no where near the kind of significant match $> 0.8$. The mean value is 0.142. These results show that the secondary fingerprint statistic clearly and reliably distinguishes between dissimilar protein structures.
